# Muscle stem cells in Duchenne muscular dystrophy exhibit molecular impairments and altered cell fate trajectories impacting regenerative capacity

**DOI:** 10.1101/2024.07.24.604963

**Authors:** Jules A. Granet, Rebecca Robertson, Alessio A. Cusmano, Romina L. Filippelli, Tim O. Lorenz, Shulei Li, Moein Yaqubi, Jo Anne Stratton, Natasha C. Chang

## Abstract

Satellite cells are muscle-resident stem cells that maintain and repair muscle. Increasing evidence supports the contributing role of satellite cells in Duchenne muscular dystrophy (DMD), a lethal degenerative muscle disease caused by loss of dystrophin. However, whether or not satellite cells exhibit dysfunction due to loss of dystrophin remains unresolved. Here, we used single cell RNA-sequencing (scRNA-seq) to determine how dystrophin deficiency impacts the satellite cell transcriptome and cellular composition by comparing satellite cells from *mdx* and the more severe D2-*mdx* DMD mouse models. DMD satellite cells were disproportionally found within myogenic progenitor clusters and a previously uncharacterized DMD enriched cluster. Despite exposure to different dystrophic environments, *mdx* and D2-*mdx* satellite cells exhibited overlapping dysregulation in gene expression and associated biological pathways. When comparing satellite stem cell versus myogenic progenitor populations, we identified unique dysfunctions between DMD and healthy satellite cells including apoptotic cell death and senescence, respectively. Pseudotime analyses revealed differences in cell fate trajectories indicating that DMD satellite cells are stalled in their differentiation capacity. *In vivo* regeneration assays confirmed that DMD satellite cells exhibit impaired myogenic gene expression and cell fate dynamics during regenerative myogenesis. These defects in differentiation capacity are accompanied by impaired senescence and autophagy dynamics. Finally, we demonstrate that inducing autophagy can rescue differentiation of DMD progenitors. Our findings provide novel molecular evidence of satellite cell dysfunction in DMD, expanding on our understanding of their role in its pathology and suggesting pathways to target and enhance their regenerative capacity.

## Introduction

Satellite cells, which are muscle-resident stem cells, are responsible for postnatal muscle growth, maintenance of muscle homeostasis, and muscle repair in response to damage^1^. Upon a regeneration-inducing stimulus, satellite cells activate and undergo myogenic commitment, transitioning into proliferative myogenic progenitors (known as myoblasts), and subsequently terminally differentiate into myocytes that undergo fusion to help repair damaged myofibers^2^. This process, known as regenerative myogenesis, is critically dependent on satellite cells^3, 4, 5^.

Recent advancement of single-cell transcriptomic technologies has allowed for the characterization of satellite cell populations from resting and regenerating muscle. These studies have not only confirmed the heterogenous nature of the satellite cell population and the myogenic trajectory of satellite cells following injury but have also revealed distinct gene expression signatures that occur due to tissue dissociation procedures^6, 7, 8, 9, 10^. Additionally, we have information pertaining to the impact of aging and muscle degenerative diseases on satellite cell heterogeneity and transcriptional changes at the single-cell level^11, 12, 13, 14, 15, 16^.

Duchenne muscular dystrophy (DMD), which impacts one in every 5,000 male births worldwide, is caused by mutations in the X-linked *DMD* gene, which lead to the loss of expression of the protein product dystrophin^17^. The vast heterogeneity of DMD mutations, the large size of the *DMD* gene, and the inefficient transduction of muscle tissue and satellite cells have impeded therapeutic development resulting in a critical need for a cure for this lethal disease. In healthy muscle, dystrophin is an integral component of the dystrophin glycoprotein complex (DGC), a large multimeric protein complex that spans the myofiber membrane, maintaining its integrity and stability^18, 19, 20^. Dystrophin is also expressed in satellite cells, where it mediates the establishment of cell polarity to enable asymmetric stem cell division^21^. Accumulating evidence supports the notion that dystrophin-deficient satellite cells exhibit intrinsic dysfunction which leads to impaired regenerative capacity^22, 23, 24^. However, whether satellite cells are directly impacted by dystrophin loss or indirectly affected due to the chronic degenerative niche, and the extent to which satellite cell dysfunction contributes to DMD pathology, has yet to be resolved. A question of particular interest is whether dystrophin-deficient satellite cells are fully capable of undergoing regenerative myogenesis and successfully contributing to muscle repair.

We sought to establish how loss of dystrophin impacts satellite cells in both moderate and severe mouse models of DMD. The *mdx* mouse is the most widely used DMD mouse model^25^. However, despite harboring a naturally occurring null mutation in the *Dmd* gene, these mice exhibit a mild dystrophic phenotype and minimal reduction in lifespan. In contrast, D2-*mdx* mice, which are *mdx* mice with a DBA/2 genetic background, exhibit a severe dystrophic pathology and a reduced lifespan of 12-18 months^26, 27^. We performed single-cell RNA-sequencing (scRNA-seq) on satellite cells enriched from *mdx*, D2-*mdx*, and their respective wildtype counterparts to assess the impact of dystrophin-deficiency on the satellite cell transcriptome and stem cell fate.

Interestingly, the majority of differentially expressed genes between the satellite cells of each DMD model and their respective controls overlapped between the two DMD models, indicating molecular impairments that are independent of the dystrophic niche. We identified distinct impairments in DMD satellite cells (apoptosis) and DMD myogenic progenitors (cellular senescence). Additionally, we characterized a DMD-specific satellite cell subpopulation that suggests an alternative, stalled differentiation fate. *In vivo* functional validation assays confirmed that DMD satellite cells exhibit impaired myogenic differentiation along with altered senescence and autophagy dynamics during regenerative myogenesis. Importantly, correcting these satellite cell dysfunctions has the potential to improve regenerative capacity, which is a critical step forward for addressing muscle repair deficits in muscle degenerative diseases.

## Results

### Altered muscle regeneration and satellite cell profiles in *mdx* and D2-*mdx* mice

To characterize muscle degeneration and the muscle-resident satellite cells in mild and severe DMD mouse models, we isolated tibialis anterior (TA) muscles from three-month-old *mdx* and D2-*mdx* mice, along with their respective wildtype counterparts C57BL/10ScSn (B10) and DBA/2 (DBA). Hematoxylin and eosin staining of TA cross-sections from *mdx* and D2-*mdx* mice revealed signs of chronic muscle degeneration (Fig. 1A) and significantly larger mean myofiber size (Fig. 1B, P = 0.0049 for *mdx* and P = 0.0217 for D2-*mdx*).

**Figure 1.**
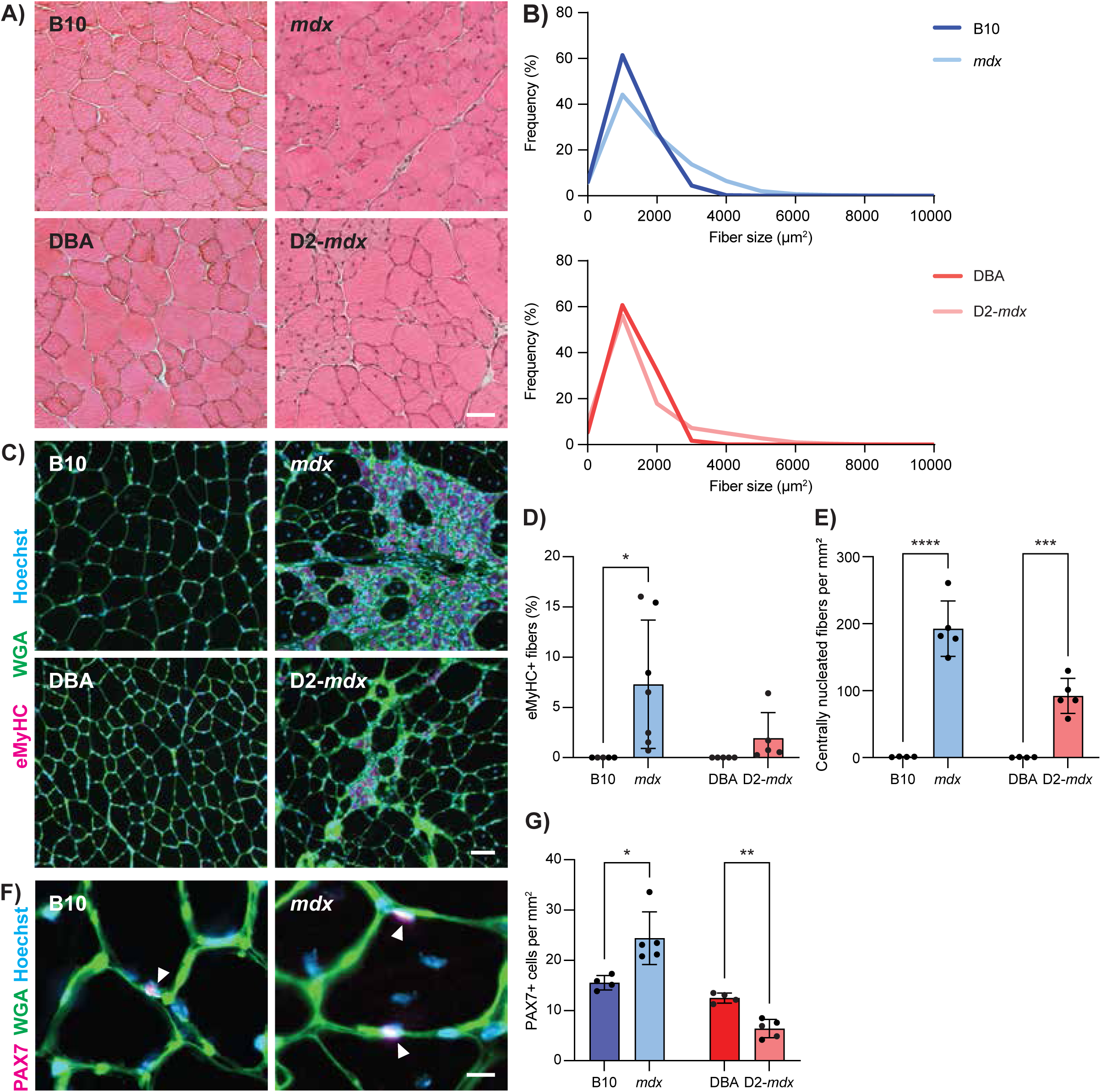
Altered regenerative and satellite cell profiles in *mdx* and D2-*mdx* mice. **A)** H&E images from TA cross-sections of three-month old male DMD models (*mdx* and D2-*mdx)* and their respective controls (B10 and DBA), showing morphological changes in the DMD models such as centralized nuclei and changes in fiber cross sectional area. **B)** Quantification of fiber cross sectional area, demonstrating changes in DMD models compared to controls. **C)** Wheat germ agglutinin (WGA, green) and eMyHC (magenta) IF labelling of TA cross-sections showing regenerating injuries only in DMD models which, as quantified in **D)** show a significant increase in eMyHC+ fibers compared to controls. **E)** Quantification of centrally nucleated fibers per mm^2^ of TA, showing a significant increase in *mdx* and D2-*mdx* compared to controls. **F)** Representative IF labelling of TA cross-sections for WGA (green) and PAX7 (magenta), showing PAX7+ satellite cells (arrows). **G)** Quantification of number of PAX7+ cells per mm^2^ of TA, showing a significant increase in *mdx* and a decrease D2-*mdx* compared to their respective controls. Scalebars = 50 µm (**A, C**) and 10 µm (**F**), *P < 0.05, **P < 0.01, ***P < 0.001, ****P < 0.0001 (two-tailed unpaired t-test, n = 4-7 biological replicates per strain). Data are expressed as frequency distribution (**B**) or mean ± SD (**D**, **E, G**).

We performed immunostaining for embryonic myosin heavy chain (eMyHC) and quantified the number of centrally nucleated myofibers on TA cross-sections to assess regeneration (Fig. 1C). While wildtype muscles exhibited little to no eMyHC+ or centrally nucleated myofibers, *mdx* and D2-*mdx* dystrophic muscles exhibited a significant increase in both the percentage of eMyHC+ myofibers and incidence of myofibers with centrally located nuclei (Fig. 1D-E). To assess satellite cell content, we performed immunostaining of TA cross-sections for the canonical satellite cell marker PAX7 (Fig. 1F). As previously reported, we found that *mdx* TA muscles have an increased number of PAX7+ satellite cells compared to B10 controls^28, 29^. In contrast, D2-*mdx* mice exhibited significantly lower numbers of PAX7+ satellite cells compared to DBA controls (Fig. 1G). When comparing *mdx* and D2-*mdx*, D2-*mdx* muscles had significantly less satellite cells compared to *mdx* (P = 0.019). Reduced numbers of satellite cells have been observed in DBA mice compared to C57BL/6 mice following repeated injuries, suggesting that differences in satellite cell content due to genetic background along with enhanced TGF-β activity contribute to the overall reduced regenerative capacity of D2-*mdx* muscles^26,30^. Consistent with this, we observe that *mdx* muscles exhibit chronic regeneration and enhanced satellite cell numbers, while D2-*mdx* muscles, despite also showing increased regeneration compared to their wildtype counterpart DBA, have significantly reduced satellite cell content and less regeneration compared to *mdx* (Fig. 1D, E, G). Altogether, these findings indicate unique differences in satellite cell and muscle regeneration phenotypes between these commonly used *mdx* and D2-*mdx* DMD models.

### scRNA-seq reveals differences in satellite cell distribution between healthy and dystrophic muscle

To focus on the satellite cell population, we performed scRNA-seq on prospectively isolated satellite cells from the hindlimb muscles of *mdx* and D2-*mdx* mice, along with their wild-type counterparts (Fig. S1A)^31^. Sequences from the four individual libraries were integrated and cell clustering was performed and visualized using uniform manifold approximation projection (UMAP) (Fig. S1B-C). Cell clusters were identified by examining highly expressed genes unique to each cluster and comparing these with published scRNA-seq data sets from satellite cells and whole muscle^7, 8, 9, 10^ (Fig. S1D). While the majority of our isolated cell population were satellite cells and myogenic progenitors (72.48-96.47%), we also detected the presence of pericyte, vascular smooth muscle, glial, Schwann, white blood, and endothelial cells (Fig. S1E-F).

Following the removal of non-myogenic and mature myonuclei cell types, sub-clustering of satellite and myogenic progenitor cells resulted in the identification of seven transcriptionally distinct clusters (Fig. 2A and S2A). Based on their gene expression signature, we designated these clusters as muscle stem cell (MuSC) 1, MuSC 2, MuSC 3, cycling progenitor, differentiating myocyte, a population comprised mainly of satellite cells from *mdx* and D2-*mdx* mice (which we termed DMD enriched), and immunomyoblast (Fig. 2B).

**Figure 2.**
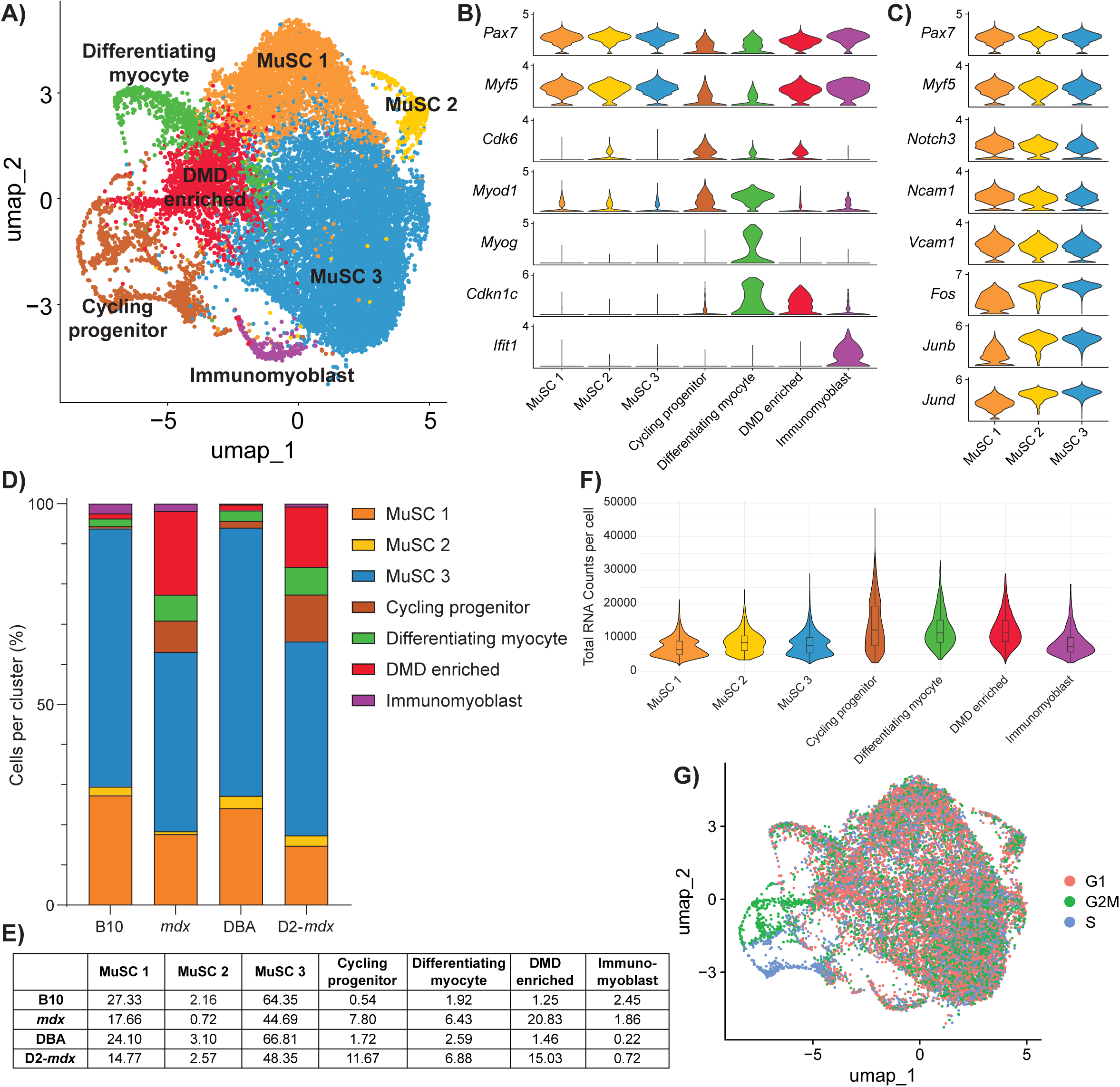
Single cell RNA-seq analysis of DMD satellite cells. **A)** UMAP representation after unsupervised clustering of myogenic cells showing the mapping of cells into seven distinct clusters. **B)** Violin plot of highly expressed markers used to define the identity of each myogenic cluster. **C)** Expanded violin plot of genes associated with satellite cell quiescence and activation for each satellite cell cluster (MuSC 1, 2 and 3). **D)** Percentage of cells belonging to each cluster by strain, summarized in **E)**, showing a decrease in MuSC clusters and increase in cycling, differentiating and “DMD enriched” clusters in the DMD models *mdx* and D2-*mdx* compared to their respective controls. **F)** Violin plot of total RNA counts per cell for each myogenic cluster. **G)** Feature plot of cell cycle stage, showing that cycling progenitors are in G2M or S phase while MuSC 1, 2, 3, differentiating myocyte, and DMD enriched cells are mainly in G1 phase.

As expected from satellite cells isolated from healthy and homeostatic resting muscle (B10 and DBA), the majority of wildtype satellite cells were found within the three MuSC clusters (93.84% in B10 and 94.01% in DBA), which express high levels of *Pax7* and *Myf5* (Fig. 2B-E). To identify where these MuSC clusters lie along the myogenic differentiation trajectory we performed gene annotation analysis on genes significantly upregulated by cells within these clusters (Fig. S2B-C). Clusters MuSC 1 and 2 uniquely expressed genes related to cell adhesion, while the MuSC 3 cluster uniquely expressed genes related to RNA transcription (Fig. S2C). Transcriptional regulatory relationships unraveled by sentence-based text-mining (TRRUST) analysis of these gene lists revealed that the MuSC 3 cluster is highly enriched for genes regulated by transcription factors associated with satellite cell activation, including Stat3, Fos, and Jun^32, 33^ (Fig. S2D). Together, these results indicate that cells within the MuSC 3 cluster exhibit higher transcriptional activity compared to MuSC 1 and 2 clusters. Moreover, among the three MuSC clusters, MuSC 1 has the lowest RNA content per cell (Fig. 2F). Based on these observations, we conclude that the MuSC clusters are representative of freshly isolated satellite cells along a continuum from close to quiescence (MuSC 1) to early activation (MuSC 3), with cells transitioning between quiescence and activation (MuSC 2). Consistent with this, cells within MuSC 1 express high levels of genes associated with satellite cell quiescence (*Notch3, Ncam1, Vcam1*)^34, 35, 36^, while cells within MuSC 3 express high levels of genes associated with satellite cell activation (*Fos*, *Junb*, *Jund*)^6, 33, 37^ (Fig. 2C).

Not surprisingly, we observed a shift in DMD satellite cells from quiescent and activated states (63.07% in *mdx*; and 65.70% in D2-*mdx*) towards myogenic progenitor states (Fig. 2D-E). These myogenic progenitor clusters included a cycling progenitor population that expresses cell cycle genes (*Cdk6*, *Mki67*) as well as the myogenic differentiation regulatory factor *Myod1*, and a differentiating myocyte population that expresses the late differentiation marker *Myog* (Fig. 2B). Cell cycle analysis confirmed that the cycling progenitor population had the highest percentage of cells in G2/M and S phase (Fig. 2G). Interestingly, a significant proportion of *mdx* and D2-*mdx* cells were found in the DMD enriched cluster (20.83% *mdx*; and 15.03% D2-*mdx*) compared to their healthy counterparts (1.25% B10 and 1.46% DBA) (Fig. 2D-E). Of note, genes that are uniquely expressed by cells in this cluster are genes associated with DMD, including *Col1a1*, *Col1a2*, *Islr*, *Mest*, *S100a11*, and *Postn*^38, 39,40, 41^. In addition to these genes, cells in this cluster also have increased expression of the negative cell cycle regulator, cyclin dependent kinase inhibitor 1C (*Cdkn1c*/*p57^kip^*^2^), which has not previously been associated with DMD (Fig. 2B, S2E). As previously reported, we also noted a minor population of immunomyoblasts (0.22-2.45%) that are enriched in immune genes (*Ifit1*, *Ifit3*, *Isg15*) (Fig. 2B, D, E)^8^.

We utilized gene expression scores from published studies and gene ontology (GO) terms to validate our cell clusters (Fig. S2F, Table S1). When we assessed the expression of these various scores across the different mouse strains, we confirmed that wildtype B10 and DBA cells are enriched for the “genuine quiescence”^42^ and “satellite cell”^43^ scores (Fig. S2G). In contrast, *mdx* and D2-*mdx* cells exhibit higher expression of the “myoblasts/myocytes”^43^ and “primed core”^6, 42^ scores. Using a DMD gene signature established from young DMD patients^38^, we also confirm that DMD satellite cells, and in particular the DMD enriched cluster are specifically expressing DMD-associated genes (Fig. S2F-G).

### DMD satellite cells exhibit overlapping dysfunctions between *mdx* and D2-*mdx* models

Principal component analysis revealed that the most significant source of variation between the four data sets is between disease (DMD) and healthy cells suggesting that loss of *Dmd* expression is driving the most significant changes between samples and not differences in the genetic background (Fig. 3A). Dystrophin-deficiency in DMD is known to impact the expression of components within the dystrophin glycoprotein complex (DGC)^44^. Thus, we assessed the expression profile of DGC genes within our scRNA-seq data set. We confirm the expression of DGC components in healthy B10 and DBA satellite cells, including dystrophin, utrophin, dystroglycan, sarcoglycans, sarcospan and syntrophin (Fig. 3B). While most DGC genes were downregulated in DMD satellite cells, *Sntb2* was an exception to this trend and exhibited increased expression (Fig. 3B).

**Figure 3.**
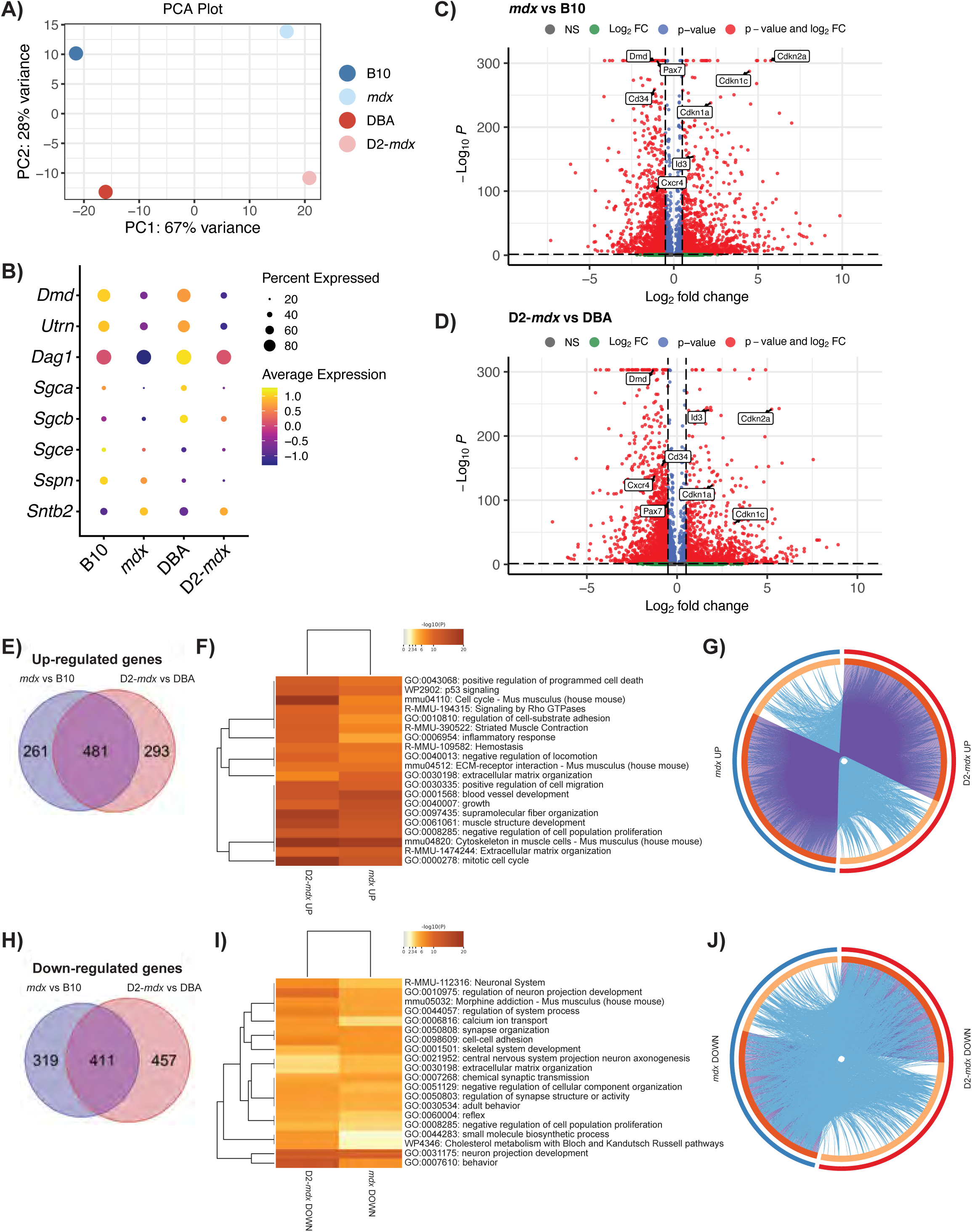
Differential gene expression in DMD satellite cells. **A)** PCA plot showing separation of DMD models versus healthy controls across PC1 and mouse genetic background on PC2. **B)** Dot plot of the expression of genes which comprise the dystrophin-glycoprotein complex, showing decreased expression of all components except *Sntb2* in DMD models vs respective controls. **C, D)** Volcano plots of differentially expressed genes in *mdx* versus B10 (**C**) and D2-*mdx* versus DBA (**D**). **E, H)** Venn diagram of up-regulated (**E**) and down-regulated (**H**) genes from *mdx* versus B10 (blue) and D2-*mdx* versus DBA (pink) and their overlap and **F, I)** heatmaps of top 20 enriched terms, coloured by p-value, related to these gene lists. **G, J)** Circos plots representing overlap between up-regulated (**G**) and down-regulated (**J**) gene lists. Outer circle represents the gene list for *mdx* (blue) and D2-*mdx* (red). Inner circle represents gene lists, where hits are arranged along the arc. Genes that hit multiple lists are colored in dark orange, and genes unique to a list are shown in light orange. Purple curves link identical genes between gene lists. Blue curves link genes that belong to the same enriched ontology term.

We performed differential gene expression analysis between DMD and healthy satellite cells (Fig. 3C-D) and between the DBA and B10 genetic backgrounds (Fig. S3A-B). We found 742 upregulated genes in *mdx* vs B10 and 774 upregulated genes in D2-*mdx* vs DBA (Fig. 3E). Surprisingly, more than half of these genes (481 genes) are overlapping between the *mdx* and D2-*mdx* sets, suggesting that despite differences in disease severity, the majority of differentially expressed genes are common to both *mdx* and D2-*mdx*. To identify biological processes related to these overlapping genes, we performed gene annotation enrichment analysis, which revealed pathways and processes including “extracellular matrix organization” (*Adamts2/4/7/14*, *Col1a1/2*, *Col5a2*, *Fn1*, *Has2*, *Mmp15*, *Tgfb1*), “cytoskeleton in muscle cells” (*Acta1*, *Myh1/3*, *Myl1*, *Tnni1*, *Tnnt2*, *Tpm2*, *Ttn*), “muscle structure development” (*Cav3*, *Cdk1*, *Id3*, *Myog*, *Sdc1*), and “mitotic cell cycle” (*Ccnb1/2*, *Cdca8*, *Cdc45*, *Cenpa*, *Cenpf*, *Cenpm*, *Kif4/11/15/20a/22/23*, *Plk1/2*) as the top enriched terms in satellite cells from both DMD models (Fig. 3F). Of note, the genes that were uniquely upregulated in either *mdx* or D2-*mdx* were still largely associated with pathways that are common to both DMD models (Fig. 3G). Altogether, these results indicate that despite disease phenotypic differences between *mdx* and D2-*mdx* models, the impact of dystrophin deficiency on satellite cells in these models are analogous.

Similarly, we observed 730 downregulated genes in *mdx* vs B10 and 868 downregulated genes in D2-*mdx* vs DBA, with 411 overlapping between the two comparisons (Fig. 3H). The top downregulated pathways included several terms associated with neural pathways including “neuron projection morphogenesis” (*Cdh4*, *Cxcr4*, *Fgfr2*, *Lgr4*, *Nlgn3*, *Ntn5*, *Vegfa*) and “cell-cell adhesion” (*Cadm2/4*, *Cdh4*, *Celsr2*, *Mcam*, *Tnxb*) (Fig. 3I-J). We noted the presence of overlapping terms between upregulated and downregulated pathways including “extracellular matrix organization” and “negative regulation of cell population proliferation” (Fig. 3F, I). While the upregulated and downregulated genes within these terms are distinct, we conclude that the presence of these terms in both comparisons indicate a general dysregulation of these pathways in DMD models.

In our comparison between the DBA and B10 backgrounds (Fig. S3A-B), “cell cycle”, “cell cycle phase transition” and “mitotic cell cycle process” pathways were upregulated in D2-*mdx* vs *mdx* (Fig. S3C-E). “Skeletal muscle cell differentiation” was downregulated in DBA vs B10, while “muscle tissue development” was downregulated in both D2-*mdx* vs *mdx* and DBA vs B10 (Fig. S3F-H). These results suggest that muscle regeneration pathways are upregulated in B10/*mdx* compared to DBA/D2-*mdx*, which is consistent with our assessment of regeneration in *mdx* and D2-*mdx* TA muscles (Fig. 1C-E)^30^.

To focus on DMD-specific gene expression differences related to cellular health that would impact myogenic differentiation, we evaluated the top 100 enriched pathways and processes and noted “positive regulation of programmed cell death” (apoptosis) and “cellular senescence”, which were increased in both DMD models compared to controls (Fig. 4A-B, S4). To determine which satellite cell clusters were most significantly impacted by apoptosis and senescence we assessed which clusters exhibited the most enrichment for these GO terms. Interestingly, the expression of genes related to apoptosis (apoptosis GO) were highest in the MuSC 2 and 3 clusters, while expression of genes related to senescence (senescence GO) was highest in the cycling progenitor cluster (Fig. 4C and S5A). These findings indicate unique cellular dysfunctions between early satellite cell and later myogenic progenitor states.

**Figure 4.**
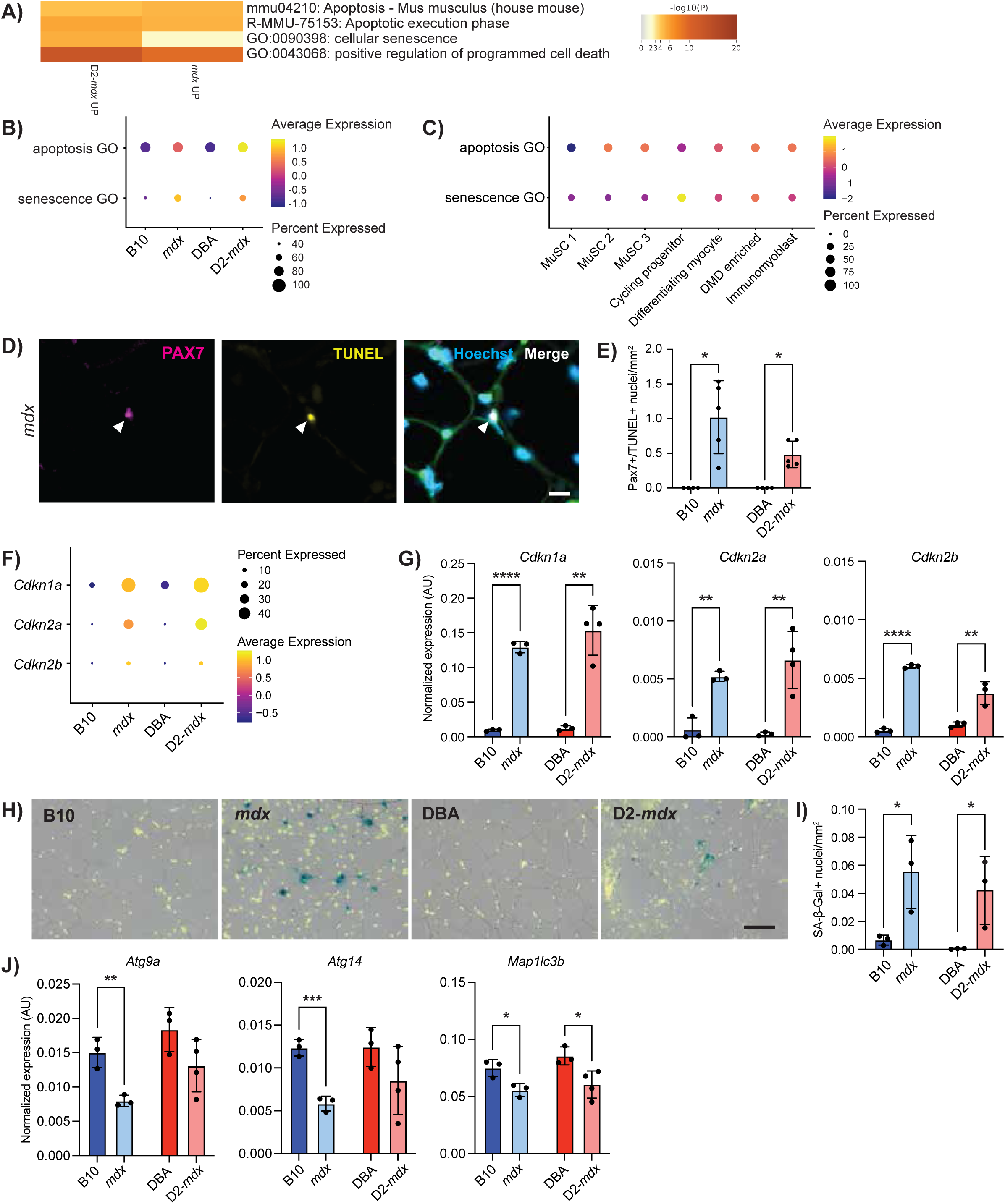
Apoptosis in DMD satellite cells and senescence in DMD myogenic progenitors. **A)** Selected terms related to apoptosis and senescence from top 100 terms derived from differentially upregulated genes in *mdx* and D2-*mdx* satellite cells compared to controls. Dot plot of the expression of apoptosis and senescence GO-terms comparing expression by **B)** strain and by **C)** cell cluster. **D)** IF labelling of a cross-section of an *mdx* TA muscle for PAX7 (magenta) and TUNEL (yellow) showing a nucleus double positive for both PAX7 and TUNEL. **E)** Quantification of number of nuclei that are PAX7 and TUNEL positive, visualized per mm^2^, showing their presence in DMD models only (n = 4-5 biological replicates). **F)** Dot plot of scRNA-seq data showing an increase in senescence-associated genes *Cdkn1a*, *Cdkn2a* and *Cdkn2b* in DMD models as compared to controls and **G)** verification of this increase in *Cdkn1a*, *Cdkn2a*, and *Cdkn2b*, with ddPCR from prospectively isolated satellite cells (n = 3 or 4 biological replicates as indicated). **H)** SA-β-Gal (blue) and Hoechst (yellow) staining of TA cross-sections showing senescence in DMD models and **I)** quantification of SA-β-Gal+ nuclei showing significant increase in DMD muscle as compared to controls. **J)** Strain-comparison of the expression of autophagy-associated genes from satellite cells described in **G**, showing reduced expression of autophagy genes in DMD models compared to healthy controls. Scalebars = 10 µm (**D**), 50 µm (**H**). *P < 0.05, **P < 0.01, ***P < 0.001, ****P < 0.0001 (two-tailed unpaired t-test (**G, I, J**), one-sided Fisher’s exact test (**E**)). Data are expressed as mean ± SD.

To examine if DMD satellite cells are indeed undergoing cell death by apoptosis, we performed terminal deoxynucleotidyl transferase dUTP nick-end labeling (TUNEL) on TA cross-sections from B10, *mdx*, DBA, and D2-*mdx* mice. In agreement with our scRNA-seq data, we observed the presence of TUNEL+ and PAX7+ satellite cells in *mdx* and D2-*mdx* muscles, indicating double stranded DNA breaks, a hallmark of apoptosis (Fig. 4D-E, S5B). In contrast, we did not observe any TUNEL+ satellite cells in B10 and DBA controls. Consistent with this data, we also performed TUNEL staining on freshly isolated myofibers from extensor digitorum longus (EDL) muscles from B10 and *mdx* mice. We observed the presence of TUNEL+ and PAX7+ satellite cells in *mdx* myofibers, which were absent in control B10 fibers (Fig. S5C-E).

DMD myogenic progenitors exhibit increased senescence, which has been reported in DMD muscles^13, 23, 45^. We assessed the expression of senescence genes *Cdkn1a* (p21^Cip1^), *Cdkn2a* (p16^Ink4a^), and *Cdkn2b* (p15^Ink4b^) in our scRNA-seq data set and confirmed increased expression and percentage of cells expressing senescence genes in DMD satellite cells compared to healthy controls (Fig. 4F). We performed quantitative droplet digital PCR (ddPCR) from freshly isolated satellite cells from B10, *mdx*, DBA, and D2-*mdx* mice and confirmed significantly increased expression of *Cdkn1a*, *Cdkn2a*, and *Cdkn2b* in both *mdx* and D2-*mdx* compared to wildtype controls (Fig. 4G). In addition, we performed senescence-associated β-galactosidase (SA-β-Gal) staining in TA cross sections from B10, *mdx*, DBA and D2-*mdx* mice, which showed significantly increased numbers of senescent nuclei per mm^2^ of tissue in DMD muscles compared to wildtype controls (Fig. 4H-I).

In healthy quiescent satellite cells, cellular autophagy, a nutrient recycling pathway, has been shown to prevent senescence and thus maintain stemness and regenerative capacity^46^. To examine if cellular autophagy is also dysregulated in DMD satellite cells we performed ddPCR analysis to assess the expression of a panel of autophagy genes (*Atg9a*, *Atg14*, *Map1lc3b*) in satellite cells freshly isolated from B10, *mdx*, DBA and D2-*mdx* mice. In agreement with studies indicating that autophagy prevents satellite cell senescence and that autophagy is dysregulated in DMD satellite cells, we found that the expression of autophagy genes were all downregulated in DMD satellite cells compared to wildtype controls^46, 47^ (Fig. 4J).

### Identification of an impaired myogenic differentiation gene signature in DMD satellite cells

Previous studies indicate that satellite cells in DMD exhibit impaired asymmetric cell division and myogenic commitment^21, 22^. We assessed the expression of components of the cell polarity complex as well as the myogenic commitment factor *Myf5* and found reduced expression of *Mark2, Prkci,* and *Myf5* in both *mdx* and D2-*mdx* satellite cells compared to controls (Fig. 5A-B). Expression of genes within the “cell polarity establishment and maintenance” GO term was also reduced in DMD satellite cells (Fig. S2G).

**Figure 5.**
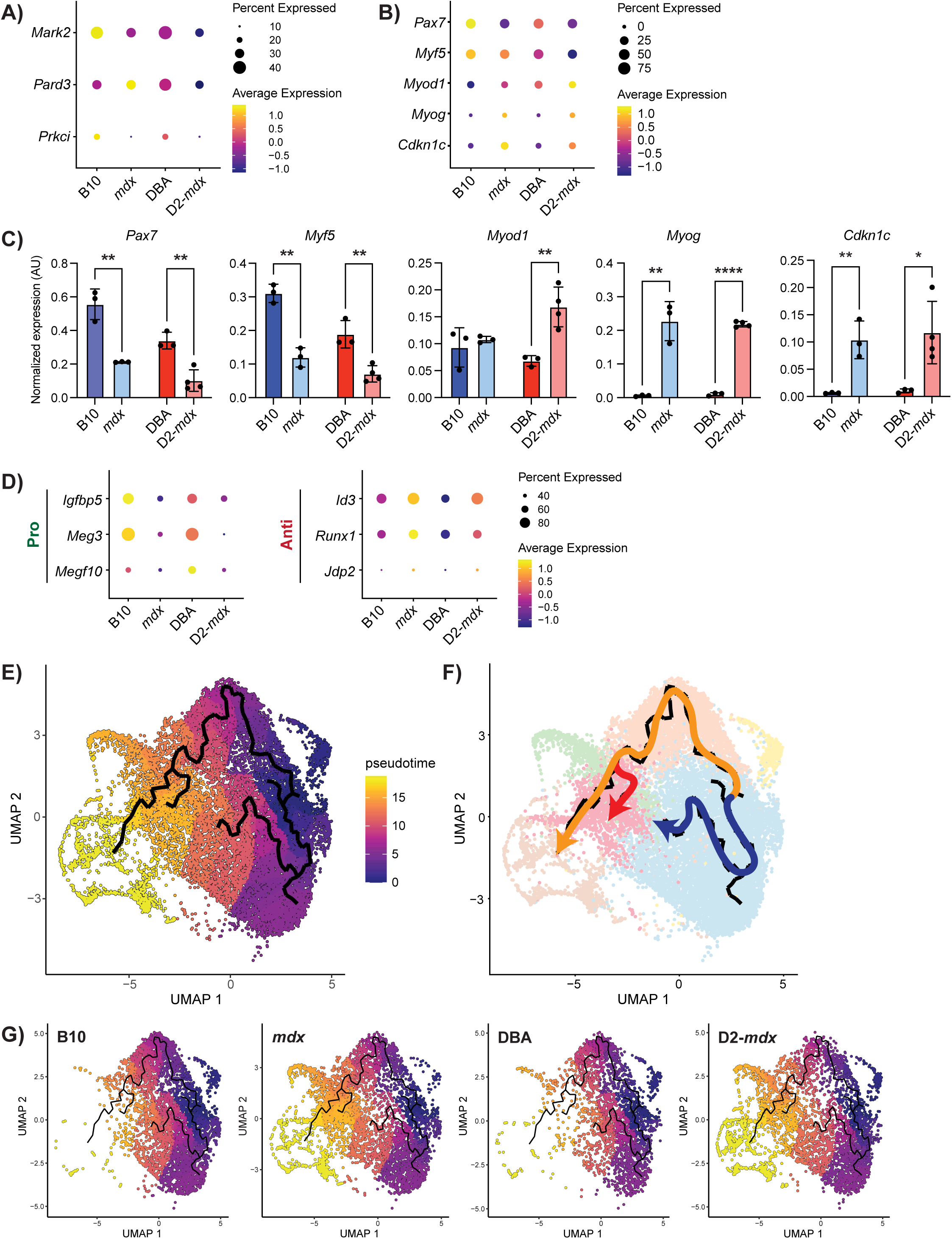
DMD MuSCs exhibit impaired myogenic differentiation. **A)** Dot plot of cell polarity genes and **B)** canonical myogenic regulatory factors and *Cdkn1c* from scRNA-seq data. **C)** Digital PCR validation of myogenic regulatory factors and *Cdkn1c* from prospectively isolated satellite cells, showing lowered expression of early satellite cell factors like *Pax7* and increased expression of late differentiation factors such as *Myog.* *P < 0.05, **P < 0.01, ****P < 0.0001 (two-tailed unpaired t-test)). Data are expressed as mean ± SD. **D)** Dot plot of scRNA-seq data for regulators of differentiation showing a decrease in pro-differentiation and an increase in anti-differentiation genes in DMD models compared to controls. **E)** Pseudotime analysis of satellite cells showing cell fate trajectories with **F)** three distinct cell fates, and **G)** visualization by strain showing a preference for the “DMD enriched” cluster fate in DMD compared to control satellite cells.

In addition to *Myf5*, we assessed expression levels of canonical satellite cell and myogenic differentiation markers in our scRNA-seq dataset (*Pax7*, *Myod1*, and *Myog*). *Pax7* and *Myf5* levels were highest in B10 and DBA cells and reduced in *mdx* and D2-*mdx* (Fig. 5B). In contrast, *Myod1* and *Myog* levels were higher in *mdx* and D2-*mdx* compared to wildtype controls. These scRNA-seq results were validated by ddPCR analysis of freshly isolated satellite cells from B10, *mdx*, DBA, and D2-*mdx* mice (Fig. 5C). The reduced expression of satellite stem cell (*Pax7*) and early myogenic commitment (*Myf5*) markers and increased expression of myogenic differentiation markers (*Myod1* and *Myog*) would suggest that DMD satellite cells exist in a more differentiated state. However, when we assessed the expression of genes involved in regulating myogenic differentiation, we found that DMD satellite cells express higher levels of anti-differentiation genes (*Id3*, *Runx1*, *Jdp2*) and lower levels of pro-differentiation genes (*Igfbp5*, *Meg3*, *Megf10*) compared to their wildtype counterparts (Fig. 5D). We also validated the increased expression of *Cdkn1c*, which coordinates the switch between proliferation and cell-cycle arrest, in freshly isolated DMD satellite cells by ddPCR^48, 49^ (Fig. 5C). Moreover, using a recently published single nuclei transcriptomic data set from muscle biopsies collected from DMD patients as well as age and gender-matched controls, we confirmed that *CDKN1C* expression is higher in DMD patient satellite cells compared to healthy controls and that expression correlates with disease severity^50^ (Fig. S5F). Together, this suggests an impairment of the normal myogenic program in DMD satellite cells.

Our findings indicate that *mdx* and D2-*mdx* satellite cells exhibit similar impairments with respect to their cellular health (apoptosis, senescence, and autophagy) and myogenic regenerative profiles. Although disease severity differs between the two DMD models, the exposure to similar dystrophic signals may contribute to the observed defects in satellite cell health and function. Thus, to assess cell autonomous defects in satellite cells prior to the dystrophic niche increasing in severity, we isolated satellite cells from prenecrotic neonate *mdx* and D2-*mdx* mice (between 10 and 14 days of age), along with their respective wildtype controls^51^. We performed gene expression using ddPCR and found that certain genes were significantly impacted in *mdx* (*Myog*, *Cdkn1c*) and D2-*mdx* (*Myf5*) neonate satellite cells compared to wildtype controls (Fig. S6A). We also observed similar trends in the senescence gene *Cdkn1a* (Fig. S6B). However, autophagy genes show opposing trends compared to adults (*Atg9a* in *mdx*) or no difference (*Atg14*) in DMD neonate satellite cells compared to control (Fig. S6C). As well, the expression of genes which are significantly impacted in both DMD strains in adults such as *Pax7* and *Cdkn2a* are not significantly changed in DMD neonate satellite cells. These findings altogether suggest that there are dysfunctions in DMD satellite cells that are cell autonomous and observed at an early stage, as well as those that are accumulated with age and exposure to a dystrophic niche.

### DMD satellite cells exhibit impaired *in vivo* regenerative myogenesis

Thus far, our analysis in homeostatic dystrophin-deficient satellite cells indicate differences in the expression of genes associated with regenerative myogenesis. To further explore this, we inferred myogenic differentiation trajectories from our scRNA-seq dataset of *mdx* and D2-*mdx* satellite cells by performing pseudotime analysis, which revealed two main trajectories (Fig. 5E-F). One trajectory, from MuSC 1 towards MuSC 3, reflects previously reported transcriptional changes that are induced by *ex vivo* muscle tissue enzymatic dissociation and satellite cell isolation procedures^6^. The other trajectory is representative of a canonical myogenic differentiation program, from MuSC 1 towards the cycling progenitor cluster, which represents the furthest pseudotime point from the MuSC clusters. The canonical trajectory included an additional branch point originating from MuSC 1 and ending in the DMD enriched cluster. Altogether, these trajectories resulted in three possible end points (MuSC 3, cycling progenitor, and DMD enriched), representing distinct cell fates. Visualization of these pseudotime trajectories on the UMAP for each mouse strain (Fig. 5G) revealed that the majority of B10 and DBA cells are found in the MuSC 3 fate. In contrast, *mdx* and D2-*mdx* cells were clearly skewed towards the cycling progenitor and DMD enriched fates.

The trajectory towards the DMD enriched cluster suggested that DMD satellite cells exhibit an altered fate, which led us to interrogate if this represents an impaired differentiate state. Genes that are highly expressed and unique to the DMD enriched cluster include several genes that have been associated with DMD as well as *Cdkn1c*. Increased expression of *Cdkn1c* in DMD satellite cells within this cluster suggests that these cells are non-proliferating due to its negative regulation of the cell cycle^48^. The DMD enriched cluster also exhibits higher *Pax7* expression and lower *Myod1* and *Myog* in comparison to the differentiating myocyte cluster (Fig. S7A), suggesting that these cells have an altered myogenic fate and are stalled in their differentiation.

To assess the differentiation potential of DMD satellite cells we utilized an *in vivo* muscle regeneration approach to induce differentiation within the context of a dystrophic niche. We performed intramuscular injury on the TA and gastrocnemius muscles of B10 and *mdx* mice and then prospectively isolated satellite cells from these mice at one- and three-days post-injury (1 and 3 DPI). Gene expression analyses by ddPCR from these samples were compared to satellite cells isolated from non-injured (NI) muscles. B10 satellite cells followed the expected trends of satellite cells transitioning from quiescence (NI) to activation (1 DPI) and proliferation (3 DPI). Expression of markers of quiescence (*Cd34*, *Calcr, Cxcr4*) as well as satellite stem cells (*Pax7*) were highest in NI cells and were reduced upon activation (Fig. 6A, S7B). *Myod1* expression was highest in B10 1 DPI activated cells, while *Myog* levels increased from 1 to 3 DPI (Fig. 6A). Interestingly, *mdx* satellite cells did not display the same dynamic expression of myogenic regulatory genes. While *Pax7* and *Myf5* expression was lower in homeostatic *mdx* cells compared to B10, expression of these factors was higher in *mdx* compared to control cells at 1 and 3 DPI. We also observed a significant reduction in *Myod1* expression at 1 DPI in *mdx* cells compared to B10. In contrast, *Myog* and *Cdkn1c* levels were elevated in *mdx* cells compared to B10 at all time points. Altogether these results indicate the dysregulation of myogenic transcription factor expression during regenerative myogenesis of DMD satellite cells.

**Figure 6.**
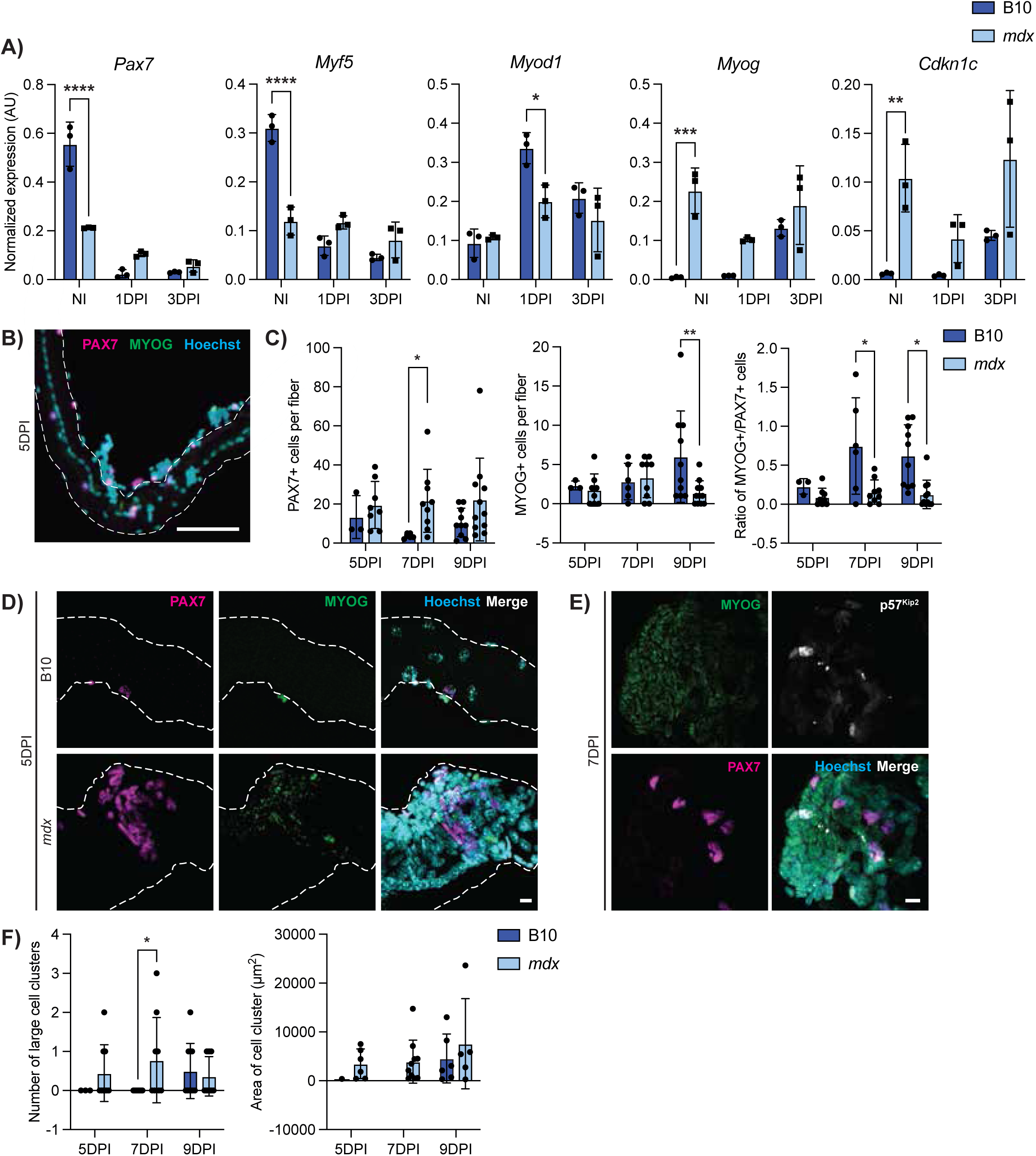
DMD satellite cells exhibit impaired *in vivo* regenerative myogenesis. **A)** Expression of myogenic factors and *Cdkn1c* in satellite cells from non-injured (NI), one-day post-injury (1DPI) and three-day post-injury (3DPI) showing dysregulation in *mdx* mice during myogenesis. (P < 0.0001 for *Pax7* and *Myf5,* P = 0.0496 for *Myod1*, P = 0.0028 for *Myog*, P = n.s. for *Cdkn1c* however strain factor P = 0.026) **B)** Representative IF labelling of an *mdx* EDL myofiber isolated five-days post-injury (5DPI) for PAX7 (magenta) and MYOG (green), showing myogenic cells along the fiber. **C)** Quantification of PAX7+ and MYOG+ nuclei at five, seven and nine DPI showing more PAX7+ and less MYOG+ nuclei in *mdx*. (Strain*DPI = n.s., strain factor P = 0.0124 for PAX7, P = 0.038 for MYOG) **D)** Images of a B10 (top) and *mdx* (bottom) EDL myofiber isolated 5DPI and labelled for PAX7 (magenta) and MYOG (green), showing a myogenic cluster in the *mdx* myofiber. **E)** A large myogenic cluster on an *mdx* EDL myofiber labelled for MYOG (green), PAX7 (magenta), and p57^Kip2^ (white) at 7DPI. **F)** Quantification of the number of large clusters (> 1000 µm^2^) and the area of these clusters. Scalebars = 100 µm (**B**), 10 µm (**D**, **E**). *P < 0.05, **P < 0.01, ***P < 0.001, ****P < 0.0001 (two-way ANOVA (**A**), two-tailed unpaired t-test (**C, F**)). Data are expressed as mean ± SD.

While ddPCR provides a quantitative readout for transcript levels, myogenic gene expression levels do not always correlate with cell fate. Thus, we also assessed myogenic protein expression and cell fate dynamics using the gold standard *ex vivo* EDL myofiber assay. Rather than isolating the myofibers and subsequently culturing them *ex vivo*, we performed intramuscular injury on the TA muscle and then collecting the adjacent EDL muscle at five-, seven-, and nine-days post injury (5, 7, and 9 DPI) to examine satellite cell differentiation that was induced *in vivo*. Quantification of myogenic cells on B10 injured myofibers revealed an expected increase in the number of MYOG+ cells from 5 to 9 DPI (Fig. 6B-C). In contrast, we did not observe an increase in MYOG+ cells in *mdx* injured fibers and the number of MYOG+ cells were significantly reduced in *mdx* fibers at 9 DPI (Fig. 6C). The number of PAX7+ cells per fiber was increased over all time points in *mdx* compared to B10 injured fibers. When representing the data as a ratio of MYOG+ to PAX7+ cells, *mdx* myofibers have significantly reduced MYOG+/PAX7+ ratios at 7 and 9 DPI. Most strikingly, we observed the presence of large clusters of cells in *mdx* fibers at all time points that were virtually absent in B10 fibers except for at 9 DPI (Fig. 6D-F). These cell clusters contained myogenic cells expressing nuclear PAX7, MYOG as well as p57^Kip2^ (protein nomenclature for *Cdkn1c*) (Fig. 6E). The numbers of large cell clusters (>1000um^2^) as well as the area of the observed cell clusters were increased in *mdx* injured myofibers compared to B10 controls (Fig. 6F). Altogether, our scRNA-seq data along with our *in vivo* regenerative myogenesis analyses of DMD satellite cells indicate that DMD satellite cells exhibit impaired myogenic differentiation.

### Altered senescence and autophagy dynamics in DMD satellite cells

Cellular senescence was enriched in DMD satellite cells compared to wildtype (Fig. 4F-I), which was specifically enriched in the cycling progenitors cluster (Fig. 4C). Thus, we assessed the expression of senescence-associated genes *Cdkn1a*, *Cdkn2a*, and *Cdkn2b* in satellite cells following injury by ddPCR. We found that in healthy B10 satellite cells, the expression of senescence genes is increased during early activation (Fig. 7A). In contrast, DMD satellite cells exhibit significantly higher baseline senescence gene expression in non-injured satellite cells and these levels are maintained during regeneration, thereby indicating a chronic persistence of senescence gene expression in DMD satellite cells, which we hypothesize is impairing the regenerative capacity of DMD satellite cells^45, 52^.

**Figure 7.**
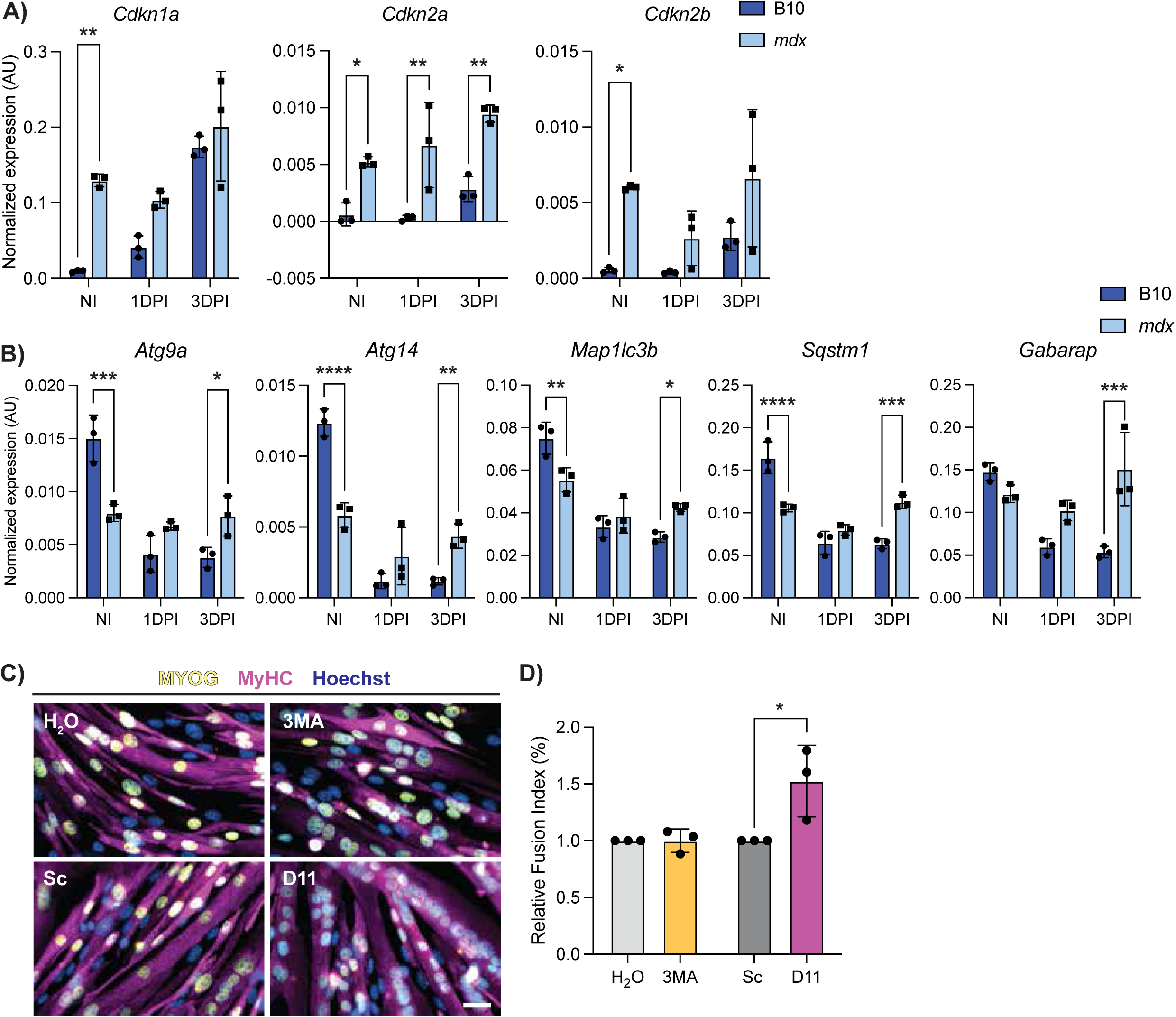
DMD satellite cells have impaired autophagy and senescence dynamics during regeneration. **A)** Digital PCR quantification of senescence-associated genes *Cdkn1a*, *Cdkn2a*, and *Cdkn2b* from non-injured (NI), one day post injury (1DPI) and three days post injury (3DPI) B10 and *mdx* mice showing dysregulation in *mdx* (P > 0.05 for effect of strain*DPI, however P < 0.0001, P = 0.0134 and P = 0.0577 for effect of strain alone). **B)** Digital PCR quantification of autophagy-associated genes in NI, 1DPI and 3DPI B10 and *mdx* satellite cells demonstrating impaired autophagy dynamics in *mdx* cells during regeneration (Strain*DPI = P < 0.0001, < 0.0001, = 0.0007, < 0.0001, and = 0.0006, respectively). **C)** Representative images of IF staining against MYOG (yellow) and MyHC (magenta) from *mdx* primary myoblasts treated with an autophagy inhibitor (3-MA, 5 mM and H_2O_ control) or autophagy inducer (Tat-D11, 10 µM and Scramble control) for two hours prior to differentiation for two days. **D)** Fusion index of each condition was determined and shows increased differentiation after Tat-D11 treatment. Scalebar = 25 µM, *P < 0.05, **P < 0.01, ***P < 0.001, ****P < 0.0001 (two-way analysis of variance (**A, B**) and two-tailed unpaired t-test (**D**)). Data are expressed as mean ± SD.

Next, we assessed the expression of autophagy genes (*Atg9a*, *Atg14*, *Map1lc3b*, *Sqstm1*, and *Gabarap*) during injury-induced myogenesis. In wildtype satellite cells, expression levels were highest during quiescence and reduced upon activation (Fig. 7B). This trend was opposite of what we observed with respect to the expression of senescence genes following injury (Fig. 7A). Moreover, this reduction in autophagy gene expression during regenerative myogenesis was absent in *mdx* satellite cells at 1 and 3 DPI suggesting impaired autophagy dynamics in *mdx* satellite cells following injury (Fig. 7B).

We assessed the expression of senescence and autophagy gene expression scores in the single nuclei human DMD transcriptomic data set^50^. Here we observed a similar trend in human DMD satellite cells as we found in our murine data from *mdx* and D2-*mdx* satellite cells. Human DMD satellite cells exhibited enhanced senescence in the stable patient group, while autophagy was reduced in both stable and declining patient groups (Fig. S7C). These data from DMD patient samples support our findings in mouse satellite cells, which are from a niche that is representative of an early disease state due to their less severe phenotype.

We next asked if modulating autophagy would improve the differentiation capacity of DMD satellite cells. To address this, we used 3-methyladenine (3MA), a class III phosphoinositide kinase inhibitor that inhibits autophagy, and Tat-Beclin 1 (D11), an autophagy promoting peptide that contains 11 amino acids from the autophagy inducing protein Beclin 1^53, 54^. We treated primary myogenic progenitors isolated from *mdx* mice with either 3MA or D11 and assessed their differentiation capacity (Fig. 7C-D, S7D-E). While inhibiting autophagy with 3MA did not have a significant impact on differentiation, inducing autophagy with D11 resulted in enhanced differentiation of DMD progenitors (Fig. 7D). Thus, inducing autophagy in DMD progenitors improves the differentiation capacity of DMD progenitor cells. Altogether, our results support the notion that counteracting these DMD progenitor-specific impairments (ie. reduced autophagy) can improve their endogenous regenerative capacity.

## Discussion

Single cell sequencing technologies have provided the ability to parse out cellular heterogeneity and identify transcriptomic differences within complex cell populations. Here, we used scRNA-sequencing to elucidate the mechanisms by which satellite cells in DMD in particular are impacted by loss of dystrophin. Recent studies have used both single cell and single nucleus transcriptomic approaches to assess whole muscle tissue from various mouse and rat DMD models^13, 14, 55^. While these approaches have identified significant alterations in tissue-wide cellular diversity in DMD, our study uniquely focused on sequencing of the satellite cell population to gain sufficient sequencing depth to identify distinct satellite cell populations within the heterogenous satellite cell pool. By adequately capturing these subpopulations, we were able to elucidate molecular pathways that differ between DMD and healthy satellite cells (apoptosis) and myogenic progenitors (senescence) and identify a DMD-specific satellite cell subpopulation.

We compared satellite cells from two DMD mouse models, the mildly dystrophic *mdx* mouse and the more severely dystrophic D2-*mdx* mouse. Of note, *mdx* and D2-*mdx* mice exhibit differences in satellite cell content. While *mdx* muscles have higher satellite cell numbers compared to their control counterpart, D2-*mdx* mice have reduced satellite cell numbers (Fig. 1G), supporting previous studies demonstrating that satellite cells have reduced regenerative capacity in DBA/2 muscles compared to C57BL/6, thereby impacting their ability to maintain a sufficient satellite cell pool within the context of chronic degeneration^26^. The altered muscle regenerative potential in DBA and D2-*mdx* is supported by our pathway enrichment analyses, which indicated reduced muscle tissue development and skeletal muscle differentiation when comparing satellite cells from the DBA versus B10 background (Fig. S3G). Whereas an increase in satellite cell numbers has been reported in human DMD tissue^28^, these differences in satellite cell number and muscle regeneration should be considered when interpreting data from the D2-*mdx* model.

Despite these differences in disease severity and thus satellite cell niche, we found the two DMD models shared many overlapping differentially expressed genes and molecular dysfunctions as compared to their healthy counterparts (Fig. 3G, J). We recognize that although *mdx* and D2-*mdx* DMD models exhibit these phenotypic differences, it is likely that dystrophic niche signaling between the two DMD models are similar. Thus, to assess dystrophin-deficient satellite cells prior to exposure to this signaling, we examined neonate satellite cells from *mdx* and D2-*mdx* mice and found early signs of myogenic impairment as well as altered senescence and autophagy gene expression (Fig. S6A-C). However, some genes, such as *Pax7* and *Cdkn2a*, do not show the same significant shift seen in adults. These findings suggest that satellite cell dysfunction in DMD is a combination of both cell autonomous defects that are present at early stages prior to muscle degeneration as well as a result of accumulated exposure to the niche. Interestingly, when we compared myogenic gene expression between neonate and adult satellite cells, the expression of myogenic genes (*Pax7*, *Myf5*, *Myog*, *Cdkn1c*) are significantly changed in adult (three months of age) vs neonate in healthy control cells (Fig. S6A). However, in DMD satellite cells this change from neonate to adult is greatly reduced, suggesting an impairment in satellite cell transition from neonate to early adult stages.

Prior to the discovery that dystrophin is expressed in the satellite cell, it was generally thought that the loss of dystrophin had minimal impact on satellite cells and their progeny, the myogenic progenitors. While earlier studied have shown no difference in or even accelerated myogenic differentiation kinetics in cultured DMD satellite cells, we propose that *in vitro* culturing selects for those cells capable of proliferating and surviving in culture rather than those vulnerable to cell death^56, 57^. There is indeed evidence that intrinsic impairment of DMD satellite cells and myogenic progenitors leads to reduced regenerative capacity, thereby resulting in an inability to support dystrophic muscle repair ^21, 22, 24, 58, 59^. Using single cell transcriptomics, we were able to visualize heterogeneity within DMD satellite cells with respect to molecular dysfunctions as well as myogenic differentiation capacity. Of note, we identified a DMD enriched cell cluster that uniquely expresses genes associated with DMD (Fig. S2F), expresses high levels of the cell cycle inhibitor *Cdkn1c* (Fig. S2E), and has a distinct cell fate trajectory (Fig. 5F). This DMD enriched cluster paradoxically expresses higher levels of stem cell and myogenic commitment markers *Pax7* and *Myf5* and lower levels of differentiation markers *Myod1* and *Myog* compared to the progenitor and myocyte clusters (Fig. S7A).

While DMD satellite cells are skewed towards progenitor populations (Fig. 2D), as expected in a continually regenerating tissue, we propose that at least a subpopulation of DMD satellite cells is stalled in differentiation and exhibit dysregulated myogenic capacity. In line with this, less than 50% of DMD satellite cells are expressing the “primed core”^42^ and “myoblast/myocyte”^43^ scores (Fig. S2G). Importantly, our *in vivo* regeneration assays confirmed an impairment in regenerative myogenesis in *mdx* satellite cells compared to B10 controls (Fig. 6A-F). Interestingly, while *mdx* satellite cells express lower levels of *Pax7* transcripts and higher levels of *Myog* transcripts, we show that the level of gene expression does not correlate with cell number as we observe increased PAX7+ cells and reduced MYOG+ cells in *mdx* fibers following injury (Fig. 6A-C). Our results suggest that *mdx* mice have a higher number of cells that are PAX7+, which each have low *Pax7* transcript, and a low number of MYOG+ cells that are expressing high levels of *Myog.* In addition, we observed previously undescribed large cell clusters uniquely in injured *mdx* myofibers (Fig. 6D-F), suggesting an impairment in cell fusion. Further experiments to isolate and characterize these cells by flow cytometry would provide important insight into this DMD enriched cell population.

We found that DMD satellite cells exhibit enrichment in programmed cell death pathways (Fig. 4A-C). This is the first reporting of increased apoptosis in DMD satellite cells. In contrast, DMD myogenic progenitors exhibit enrichment in cellular senescence (Fig. 4A-C). We found that healthy satellite cells express increased levels of senescence-associated genes following injury (Fig. 7A). Recent studies have indicated that senescent cells are a normal component within the regenerating muscle niche^11, 16, 60^, however DMD satellite cells exhibit increased baseline expression of senescence-associated genes suggesting a chronic and dysregulated senescent state (Fig. 4G, 7A). The chronic presence of these senescent cells in DMD would hinder muscle regeneration^16, 45^.

The nutrient recycling macroautophagy pathway has previously been shown to be dysregulated in DMD muscle and satellite cells^47, 61, 62^. Consistent with these findings, we found that the expression of autophagy genes is universally lower in DMD satellite cells (Fig. 4J) and do not exhibit a dynamic downregulation during regenerative myogenesis (Fig. 7B). We propose that this impairment in autophagy dynamics would also impact the regenerative capacity of DMD satellite cells. Indeed, we found that stimulating autophagy with Tat-Beclin D11, which induces autophagy independently of inhibiting the master metabolic regulator mammalian target of rapamycin, improved the differentiation capacity of *mdx* progenitor cells (Fig. 7C-D)^53^. Our results support earlier work indicating that autophagy reduction in DMD muscle tissue correlates with reduced muscle function and that inducing autophagy with agents such as rapamycin and the AMPK agonist AICAR can improve regeneration and pathology in *mdx* mice^47, 63, 64^.

In conclusion, our study provides additional molecular evidence that satellite cells are dysfunctional in DMD and that their reduced regenerative potential contributes to the characteristic progressive muscle wasting of the disease. The work described here serves as a critical resource characterizing the transcriptomes of two widely used DMD mouse models, *mdx* and D2-*mdx*, along with their wildtype counterparts. Importantly, transcriptomic data from satellite cells from DMD patients supports our findings of dysregulated cell death, senescence, and autophagy pathways. Our results indicate that DMD satellite cells are heterogenous with respect to their myogenic regenerative potential and their exhibition of impaired cellular pathways. We propose that gene therapies aimed at transducing satellite cells should examine the benefits and consequences of targeting specific satellite cell populations. Overall, our study provides a molecular map that illustrates how satellite cell subpopulations are impacted by the loss of dystrophin and will inform therapeutic strategies aimed at improving the regenerative capacity of DMD muscle.

## Methods

### Mice

Housing, husbandry and all experimental protocols for mice used in this study were performed in accordance with the guidelines established by the McGill University Animal Care Committee, which is based on the guidelines of the Canadian Council on Animal Care. C57BL/10ScSnJ (B10, #000476), C57BL/10ScSn-*Dmd^mdx^*/J (*mdx*, #001801), DBA/2J (DBA2, #000671), and D2.B10-*Dmd^mdx^*/J (D2-*mdx*, #013141) were purchased from The Jackson Laboratory. One- to one-and-a-half-week-old male mice were used for neonatal experiments and two- to three-month-old male mice were used for experiments in adults.

### Muscle histology

Cryostat cross-sections of tibialis anterior (TA) muscle of 10 μm were used for hematoxylin and eosin (H&E) staining, senescence-associated β-galactosidase staining and immunofluorescent (IF) labelling. Sections for H&E were fixed with 4% paraformaldehyde (PFA) and stained with Mayer’s modified hematoxylin solution (Abcam) and washed for five minutes followed by differentiation with 1% HCl in 10% ethanol. Sections were stained with Eosin Y working solution (0.25% Eosin disodium salt (Sigma) in 80% ethanol with 0.5% glacial acetic acid), dehydrated and mounted with xylene mounting media (Fisher).

For terminal deoxynucleotidyl transferase dUTP nick end labelling (TUNEL)/PAX7 immunofluorescence (IF) co-labelling, sections were fixed with 2% PFA for ten minutes, then decrosslinked for 45 minutes at 95 °C in citrate buffer (Abcam). This was followed by permeabilization and blocking for one hour (0.25% triton X-100 + 0.1 M Glycine in PBS + 5% donkey serum (DS) + 2% bovine serum albumin (BSA) + Mouse on Mouse Blocking Reagent (Vector Laboratories) in phosphate buffered saline (PBS)). TUNEL assay was done by incubating slides with an *In Situ* Cell Death Detection Kit (Roche) for 37 °C for half an hour. Primary antibody incubation was performed overnight at 4 °C with pure PAX7 antibody (DSHB) with 5% DS + 2% BSA. Secondary incubation in labelling buffer (5% DS + 2% BSA in PBS) was done for one hour at room temperature, followed by Hoechst staining for five minutes. Slides were mounted with ProLong^TM^ Gold antifade reagent (Thermo Fisher).

For eMyHC labelling, TA cryosections were fixed with 4% PFA for 10 minutes then permeabilized and blocked for one hour (0.1 M glycine + 5% DS + 0.25% triton-X-100 + Mouse on Mouse Blocking Reagent (Vector Laboratories) in tris buffered saline (TBS)). Subsequently, cryosections were incubated overnight at 4 °C with eMyHC primary antibody (DSHB) primary antibody solution (1% DS + 0.25% triton-X-100 in TBS). Secondary antibody incubation and mounting were performed as above.

Sections were imaged using a Zeiss Axio Observer 7 at 20X, except for eMyHC-stained cryosections which were imaged at an EVOS M5000 microscope using a 10X objective.

### Senescence-associated β-galactosidase staining

Tibialis anterior (TA) sections were fixed for four minutes with 1% PFA and 0.2% glutaraldehyde in PBS, followed by incubating for 30 minutes in PBS at pH 5.5. Sections were then incubated with X-gal staining solution (4 mM potassium ferricyanide + 4 mM potassium ferrocyanide + 2 mM MgCl_2 +_1 mg/mL X-gal and 0.04% IGEPAL (Sigma) in PBS at pH 5.5) for 48 hours at 37 °C. Sections were subsequently washed in PBS for 10 minutes and fixed again with 1% PFA diluted in PBS for 10 minutes. Sections were counterstained with Hoechst. The stained sections were mounted with ProLong^TM^ Gold antifade reagent and imaged using a Zeiss Axio Observer microscope.

### Single myofiber isolation

Single myofibers were isolated from the extensor digitorum longus (EDL) muscle of mice that were either uninjured or mice that received injections of 1.2% BaCl_2 t_o the TA (30 µl at 1.2%) five, seven or nine days prior to isolation to induce injury and thus activate satellite cells^65^. After isolation, single myofibers were collected and fixed immediately in 2% PFA.

### IF and TUNEL staining on fibers

Fixed myofibers were permeabilized for 10 minutes (0.1 M glycine + 0.1% triton X-100 in PBS) then blocked for two hours (2% BSA + 2.5% DS + 2.5% goat serum (GS) in PBS). Primary antibody incubation was done overnight labelling solution (0.5% DS + 0.5% GS in PBS) at 4 °C with antibodies for PAX7 (DSHB), MYOG (Abcam), p57^Kip2^ (Santa Cruz). Secondary antibody incubation was done for one hour at room temperature in labelling solution, followed by Hoechst staining for one minute. Fibers were mounted on slides with ProLong^TM^ Gold antifade reagent.

TUNEL staining was done with the *In Situ* Cell Death Detection Kit (Roche). Fibers were fixed with 2% PFA, then permeabilized and blocked as previously described. Fibers were then incubated with TUNEL reaction mixture for one hour at 37 °C followed by primary antibody incubation overnight at 4 °C. Secondary antibody incubation, Hoechst and mounting on slides were performed as described above.

Myofibers were imaged using a Zeiss LSM710 using ZEN 3.2 (blue edition), with the exception of Fig. 6B which was imaged at an EVOS M5000 microscope (software version 1.6.1899.478) using a 20X objective.

### Quantification of IF images

All image analysis was done with Fiji ImageJ (version 1.45f)^66^. PAX7+, MYOG+, TUNEL+, and SA- Βgal+ nuclei quantification was performed using the cell counter feature. Cross sectional area of myofibers and aspect ratio was obtained using the automated muscle histology analysis algorithm Myosoft (version 14) to define individual fiber ROIs via WGA staining^67^. Using this same software, the number of eMyHC+ fibers were determined by automatically measuring mean grey value. These values were then corrected through background subtraction. A negative control was used to determine the range of values that represented “true” negatives and values above this (>75) were considered eMyHC+. Fibers that contained one or more nuclei within the fiber were counted as centrally nucleated fibers.

### Satellite cell isolation

Satellite cells were prospectively isolated from the hind limb muscles of mice by fluorescence-activated cell sorting (FACS) as previously described^68^. Cells were labelled with negative and positive lineage markers (Table S2) and satellite cells were sorted using a BD FACSAriaTM III (BD Biosciences).

### Mouse single cell RNA-sequencing and computational analyses

For each mouse strain (B10, *mdx*, DBA, D2-*mdx*), ITGA7+/VCAM+/Lin-satellite cells from two mice were pooled, sorted and captured on the 10X Genomics Chromium platform and subjected to single cell transcriptomic sequencing. Following satellite cell isolation, the single-cell RNA-sequencing libraries were prepared using the 10X Genomics NextGen scRNA 3’ V3.1 (10X Genomics, Pleasanton, CA) with approximately 10,000 satellite cells from each strain. Libraries were then sequenced with an Illumina NovaSeq6000 sequencer (Illumina, San Diego, CA). Reads were processed with Cell Ranger (version 3.0.1) and aligned to the mouse reference transcriptome mm10.

Computational analysis was performed using RStudio (version 4.2.1, 2022-06-23), and the data were imported with Seurat (version 5.1.0)^69^. Cells were filtered to remove cells with less than 1,000 or more than 5,000 genes, as well as cells with more than 8% mitochondrial genes, to eliminate low quality cells. Doublets were removed using DoubletFinder (version 2.0.4)^70^. After normalization and scaling, cells from the different mouse strains were integrated. The resolution was determined using the Clustree function (version 0.5.1) and unsupervised clustering was carried out using the FindClusters function^71^. The first clustering was done with a resolution of 0.1, and the sub-clustering of the myogenic cells (MuSC and myoblast) was done with a resolution of 0.2. Cells expressing high levels mature myonuclei genes (*Acta1*, *Tnnc2*, *Tnnt3*), representing a minor portion of total myogenic cells (0.04 – 0.11%) were removed for final downstream analysis. Differential gene expression and cluster identification were performed using FindAllMarkers function in Seurat. The scores were generated with the AddModuleScore function, using published genes lists. Stacked violin plots were made using the StackedVlnPlot function^72^. Dot plots and feature plots were created using scCustomize package (version 2.1.2)^73^. Cell cycle analysis was done using the CellCycleScoring function of the Seurat Package. The total RNA counts per cell were done by extracting the raw count matrix using the GetAssayData function, and the total RNA counts (UMI) per cell were calculated and added to the metadata. The violin plot was created using ggplot2 package (version 3.5.1).

### Human single nuclei RNA-sequencing and computational analyses

Control and DMD human data were generated and kindly shared by Prof. Jordi Diaz-Manera’s team^50^. Computational analysis was performed as described above. Cells were filtered to remove cells with less than 100 or more than 5,000 genes, as well as cells with more than 5% mitochondrial genes, and doublets were removed using DoubletFinder (version 2.0.4)^70^. After normalization and scaling, cells from the different samples were integrated with a resolution of 0.1, and the sub-clustering of the myogenic cells (*PAX7*^+^) was done with a resolution of 0.4. The sample groups were generated as described^50^.

### Principal component analysis (PCA) – Pseudobulk

The processed individual datasets were merged, and the object was converted into a SingleCellExperiment object. Counts were aggregated with aggregateBioVar package (version 1.14.0)^74^. Differential gene expression was performed using DESeq2 Package (version 1.44.0)^75^. The PCA plot was generated using plotPCA function.

### Differential gene expression and Gene Ontology (GO) enrichment analysis

Differentially expressed genes lists were generated using the FindMarkers function. Genes with Log2FC more than 1 and adj pVal less than 0.05 were selected for GO enrichment analysis. GO enrichment was performed using Metascape (version 3.5, metascape.org) with multiple gene lists using the Custom Analysis function^76^. GO Biological Processes, KEGG Pathway, Reactome Gene Sets, CORUM, WikiPathways, and PANTHER Pathway were selected. Terms with a p-value < 0.01, a minimum count of 3, and an enrichment factor > 1.5 were grouped into clusters based on their membership similarities. GO terms analysis has been performed with the enrichGO function of the clusterProfiler package version 4.12.0^77^. Volcano plots were generated using the EnhancedVolcano package (version 1.22.0)^78^. Venn Diagrams were generated using VennDiagram package (version 1.7.3)^79^.

### Pseudotime analysis

The pseudotime analysis was performed using Monocle3 package (version 1.3.7)^80^. The datasets were processed as described previously. The Seurat object has been transformed as a cell data set using the SeuratWrappers package (version 0.3.5). Cells were then clustered using the cluster_cells function. The trajectory was generated using the learn_graph function by adjusting the default setting of minimal_branch_len to 10, euclidean_distance_ratio to 1 and geodesic_distance_ratio to 1. Finally, cells were ordered along the learned trajectory.

### Droplet digital PCR (ddPCR)

RNA was extracted from isolated satellite cells using the PicoPure RNA Isolation Kit (Applied Biosystems) and cDNA was generated with the SuperScript III First-Strand Synthesis System (Invitrogen). ddPCR was performed using ddPCR Supermix for Probes (no dUTP) (Biorad) and data collected with the QX100 Droplet Digital PCR System (Biorad) with QuantaSoft (version 1.7.4). A list of primers can be found in Table S3.

### Autophagy modulation assays

Primary satellite cell-derived myoblasts were isolated from *mdx* mice as previously described^81^. Cells were seeded in proliferation media (20% FBS, 10% HS, 3% CEE, 10 ng/mL bFGF and 2 mM L-glutamine in Ham’s F-10) on collagen-coated 35 mm plates. When cells reached 80-90% confluence, cells were treated for two hours with either 10 µM Tat-Beclin 1 D11 peptide (Novus Biologicals) or Tat-Beclin 1 L11S peptide scramble control (vehicle, Novus Biologicals); 5 mM 3-methyladenine (3-MA, Sigma) or water (vehicle)^53, 54^. Following treatment, cells were induced to differentiate for two days in differentiation medium (50% DMEM, 50% F10, 5% HS). Cells were fixed and immunolabeled with MYOG (Abcam) and MyHC antibody (DSHB), and fusion index was determined as previously described^82^.

### Western blotting

Cell pellets from a replicate of the autophagy modulation assays were collected in lysis buffer (50 mM Tris pH7.5, 150 mM NaCl, 2 mM MgCl2, 0.5 mM EDTA, 0.5% Triton X-100, 1X protease inhibitor, and 1X phosphatase inhibitor) and incubated on ice for 30 minutes. Samples were centrifuged at 15,000 rcf at 4°C for 20 minutes and supernatants collected. The protein concentration of the resulting lysates was determined using the Pierce BCA Assay Kit (Thermo Fisher). Lysates were mixed with 4X Laemmli sample buffer, denatured at 95°C for five minutes, and resolved using an 12% SDS-PAGE gel containing 0.5% 2,2,2-Trichloroethanol (TCE). Total protein was determined using the ChemiDoc imaging system (Bio-Rad) via ultra-violet activation. Samples were then transferred to a polyvinylidene difluoride (PVDF) membrane and blocked for one hour at room temperature in blocking buffer (5% milk + 0.5% Tween in PBS). Membranes were incubated with primary antibody in blocking buffer at 4°C overnight with agitation, then incubated with secondary antibody in blocking buffer for one hour at room temperature. Membrane was incubated with SuperSignal West Femto Maximum Sensitivity Substrate (Thermo Fisher) and visualized using the ChemiDoc imaging system (Bio-Rad). Protein band intensity was quantified using Fiji ImageJ.

### Statistical analysis

For data not related to scRNA-seq, statistical analyses and visualizations were done in GraphPad Prism (version 10.2.2). Significance was determined through unpaired t-tests, two-way analysis of variance (ANOVA) with Šídák’s multiple comparisons test, mixed-effects analysis with Šídák’s multiple comparisons test, or one-sided Fisher’s exact test, as indicated. P values in figure legends for two-way ANOVAs and mixed-effects analysis describe row*column effect unless otherwise noted. Confidence interval = 95% for all tests.

Two-tailed t-tests in Fig. 1 were performed on 4-7 biological replicates per group. Analysis of fiber area was done via nested t-test. Analysis of PAX7+/TUNEL+ satellite cells (Fig. 4D-E) was done with a one-sided t-test as control animals are assumed to have few such cells. For all two-tailed unpaired t-tests of strain comparison ddPCR data, n = 3 for all groups except D2-*mdx* where n = 4. Two-way ANOVA tests which were conducted on NI, 1 DPI and 3 DPI data and data comparing neonates to adults. For injury data, three biological replicates were used for all groups except B10 3DPI, where n = 4. For neonate data, n = 3-5 biological replicates as indicated on graphs. Cluster data in Fig. 6C and F was done using EDL fibers from one biological replicate per condition. Significance was then calculated via mixed-effects analysis due to uneven sample size. β-galactosidase assays were done with three biological replicates per condition and significance calculated via two-tailed unpaired t-tests. TUNEL+ nuclei percentage from EDL fibers was calculated from three mice per strain, counting n = 25, 50 and 21 B10 and n = 82, 45 and 69 *mdx* PAX7+ nuclei. A one-sided Fisher’s exact test was used as B10 has zero positives. Autophagy modulation assays were done on three biological replicates per condition and significance determined with two-tailed unpaired t-tests. Standard sample sizes for the field and that are sufficient to show differences with this disease model were chosen. scRNA-seq sample size was chosen to ensure sufficient depth of sequencing.

## Data availability

Integrated scRNA-seq datasets generated in this study are available at: https://singlocell.openscience.mcgill.ca/display?dataset=Muscle_Stem_Cells_DMD_2024

## Acknowledgements

We thank Camille Stegen and Julien Leconte from the Flow Cytometry Facility, Frederique Dansereau and Giovanni Carrera-Viray from the Comparative Medicine and Animal Resources Centre, Tony Kwan and Ashot Harutyunyan from the McGill Genome Centre, and Alexis Allot from the Single Cell Neurobiology Hub at McGill University. We also thank Prof. Jordi Diaz-Manera for kindly sharing DMD patient transcriptomic data. We are grateful to present and past members of the lab, Mitchell Wong, Julie Thomas, Lee Rowsell, and Gurtaaj Gill for their contributions. J.A.G. is the recipient of a Herbert A. Davis, M. Eve Cameron-Davis and Derek H. Davis Fellowship from McGill University. R.R. was a Muscular Dystrophy Canada Fellow. A.A.C. is the recipient of a Canada Graduate Scholarship - Master’s award. R.L.F. is the recipient of a Vanier Award (186914). S.L. is supported by the Défi-Canderel studentship from the Rosalind and Morris Goodman Cancer Institute and a Master’s training scholarship from the Fonds de recherche du Québec – Santé (328456). N.C.C. is the recipient of a Chercheur-boursier Junior 1 award from the Fonds de recherche du Québec – Santé (298146 and 309857). These studies were carried out with grants to N.C.C. from the McGill Regenerative Medicine Network, Stem Cell Network (ECR-C4R1-2), The Jesse Davidson Foundation - Defeat Duchenne Canada, and the Canadian Institutes of Health Research (175431 and 180499). We also acknowledge the support from the Neuromuscular Disease Network for Canada (NMD4C) and Quebec Cell, Tissue and Gene Therapy Network - ThéCell.

## Author contributions

N.C.C. conceived and managed the project. J.A.G., R.R., A.A.C., R.L.F., T.O.L., S.L. designed and performed experiments, and collected data. J.A.G., R.R., A.A.C., R.L.F., and N.C.C. analyzed and interpreted the data. M.Y. and J.A.S. contributed bioinformatics analysis. N.C.C. and R.R. wrote the manuscript. All authors carefully reviewed and provided critical insights into the manuscript.

## Competing interests

The authors declare no competing interests.

## Notes

### Competing Interest Statement

The authors have declared no competing interest.

### Summary of Updates

Additional replicates and experiments added to Figure 1; Single cell analysis updated to clarify differences between muscle stem cell clusters (Figure 2 revised); Figures 4-7 revised; Supplemental files updated.

