## Supplemental Information for "Muscle stem cells in Duchenne muscular dystrophy exhibit molecular impairments and altered cell fate trajectories impacting regenerative capacity"

**Supplementary tables:**

**Table S1.** Gene lists from published studies or the Gene Ontology knowledgebase used to generate the scores (Figures 4B-C, S2F-G, S5A, and S7C).

**Table S2.** Antibodies used and applications.

**Table S3.** Primer and probe sequences used in ddPCR experiments.

**Table S2**. Antibodies used and applications.

| Application | Antibody | Dilution |
|---|---|---|
| Primary antibody (IF) | Anti-Pax7 (DSHB PAX7) | None |
| | Anti-MYOG (Abcam ab124800) | 1:200 |
| | Anti-eMyHC (DSHB F1.652) | None |
| | Anti-MyHC (DSHB MF20) | 1:3 |
| | Anti-p57$^{Kip2}$ (Santa Cruz sc-563410) | 1:50 |
| Secondary antibody (IF) | Donkey anti-Mouse IgG (H+L), Alexa 647, A3278 | 1:1,000 |
| | Donkey anti-Rabbit IgG (H+L), Alexa 647, A32795 | 1:1,000 |
| | Goat anti-Mouse IgG1 (y1), Alexa Fluor 647, A21240 | 1:1,000 |
| | Goat anti-Mouse IgG2b (y2b), Alexa Fluor 555, A21147 | 1:1,000 |
| | Wheat germ agglutinin (WGA) (Life Technologies, W11261) | 1:100 |
| Primary antibody (western blot) | Anti-LC3B (Novus NB100-2220) | 1:3,000 |
| | Anti-p62 (Sigma P0067) | 1:4,000 |
| Secondary antibody (western blot) | Goat Anti-Rabbit IgG (H + L)-HRP Conjugate (Bio Rad 1706515) | 1:10,000 |
| Positive selection (FACS) | a7-integrin-Alexa 647 (UBC Ablab 67-0010-05) | 1:1,000 |
| | VCAM-PE/Cy7 (BioLegend 105720) | 1:400 |
| Negative selection (FACS) | CD31-PE (BD Biosciences 553373) | 1:10,000 |
| | CD45-PE (BD Biosciences 553081) | 1:10,000 |
| | CD11b-PE (BD Biosciences 553311) | 1:10,000 |
| | Ly6A/E (Sca-1)-PE (BD Biosciences 553108) | 1:10,000 |

**Table S3.** Primer and probe sequences used in ddPCR experiments. PrimeTime™ qPCR Probe Assays or custom assays were purchased from Integrated DNA Technologies (IDT).

| **Target** | **IDT Assay ID** | **Primers** | **Probe** |
|---|---|---|---|
| *Atg9a* | Mm.PT.58.13116605 | F: CGAGGCTGGTAACTGGAATC<br>R: CCTGTCCACCTTGTTAACCA | CTGAGCCCCAACT<br>GTCCAACCA |
| *Atg14* | Mm.PT.58.12027737 | F: GCGGTGATTTCGTCTATTTCG<br>R: GCTTGTTCTTAAGTTGGCTTAGTC | TGTCAATAAACCT<br>CTCCCGGTCGC |
| *Calcr* | Mm.PT.58.13013231 | F: GAAGATCAGCATGGAAGCAAC<br>R: TCCAACTATACTCTGTGCAACG | CAGCAATCGACAAGGA<br>GTGACCCA |
| *Cd34* | Mm.PT.58.8626728 | F: CGTGGTAGCAGAAGTCAAGT<br>R: GGTACAGGAGAATGCAGGTC | ATGCAGCAGACTCATC<br>AGGCAGAG |
| *Cdkn1a* | Mm.PT.58.17125846 | F: GAAGAGACAACGGCACACT<br>R: CAGATCCACAGCGATATCCAG | TTCAGAGCCACAGGCA<br>CCATGT |
| *Cdkn1c* | Mm.PT.58.13296375 | F: GCAGTTCTCTTGCGCTTG<br>R: CAGGACGAGAATCAAGAGCAG | AAGAAGTCGTTCGCAT<br>TGGCCG |
| *Cdkn2a* | Mm.PT.58.8388138 | F: AATCTGCACCGTAGTTGAGC<br>R: TGTTGTTGAGGCTAGAGAGGA | CGCACCGGAATCCTGG<br>ACCA |
| *Cdkn2b* | Mm.PT.58.7138437 | F: GTGCACAGGTCTGGTAAGG<br>R: AGATCCCAACGCCCTGAA | TCCACGGAGCAGAACC<br>CAACTG |
| *Cxcr4* | Mm.PT.58.41597935 | F: CCCACTTCTTCAGAGTAGTTATCAG<br>R: CGTTTGGTGCTCCGGTAA | TGCCATGGAACCGATC<br>AGTGTGAG |
| *Gabarap* | Mm.PT.58.10894755 | F: ACAGACCATAGACGCTTTCATC<br>R: CGTGCTGAAGATGCCTTGT | CAATGTCATTCCACCCA<br>CCAGTGC |
| *Map1lc3b* | Mm.PT.58.29764292 | F: GAAGATGTCCGGCTCATCC<br>R: CCAGGAACTTGGTCTTGTCC | TGCTTCTCCC<br>CCTTGTATCGCTCTAT |
| *Myf5* | Mm.PT.58.5271235 | F: ACATGCATTTGATACATCAGGAC<br>R: CACCTCCAACTGCTCTGAC | TGCCTGAATGTAA<br>CAGCCCTGTCTG |
| *Myod1* | Custom | F: ACTACAGTGGCGACTCAGAT<br>R: TGTAGTAGGCGGTGTCGTAG | TCTGATGGCATGATGG<br>ATTACAGCGG |
| *Myog* | Mm.PT.58.6732917 | F: GACCGAACTCCAGTGCATT<br>R: CTTGCTCAGCTCCCTCAAC | AGCCCATGGTGCC<br>CAGTGAAT |
| *Pax7* | Mm.PT.58.9286978 | F: TGTGACGGATGTGGTTCG<br>R: TCCCCAGGATGATGAGACC | TTGATGAAGACCCCAC<br>CAAGCTGAT |
| *Rps18* | Mm.PT.58.12109666 | F: ACACCACATGAGCATATCTCC<br>R: CCTGAGAAGTTCCAGCACAT | AGCCTTCGCCATCACTG<br>CCATTA |
| *Rps20* | Mm.PT.58.41623895.g | F: CGATTTATTCAGTTGTCTTAGGCATC<br>R: AGTCCTTCTGAGATTGTTAAGCAG | CGGGAGTTGAGGTTGA<br>AGTCACCA |
| *Sqstm1* | Mm.PT.58.5854953 | F: TGTCGTAATTCTTGGTCTGTAGG<br>R: TGCCCTATACCCACATCTCC | TGAGTCCCTCTCCCAGA<br>TGCTGT |

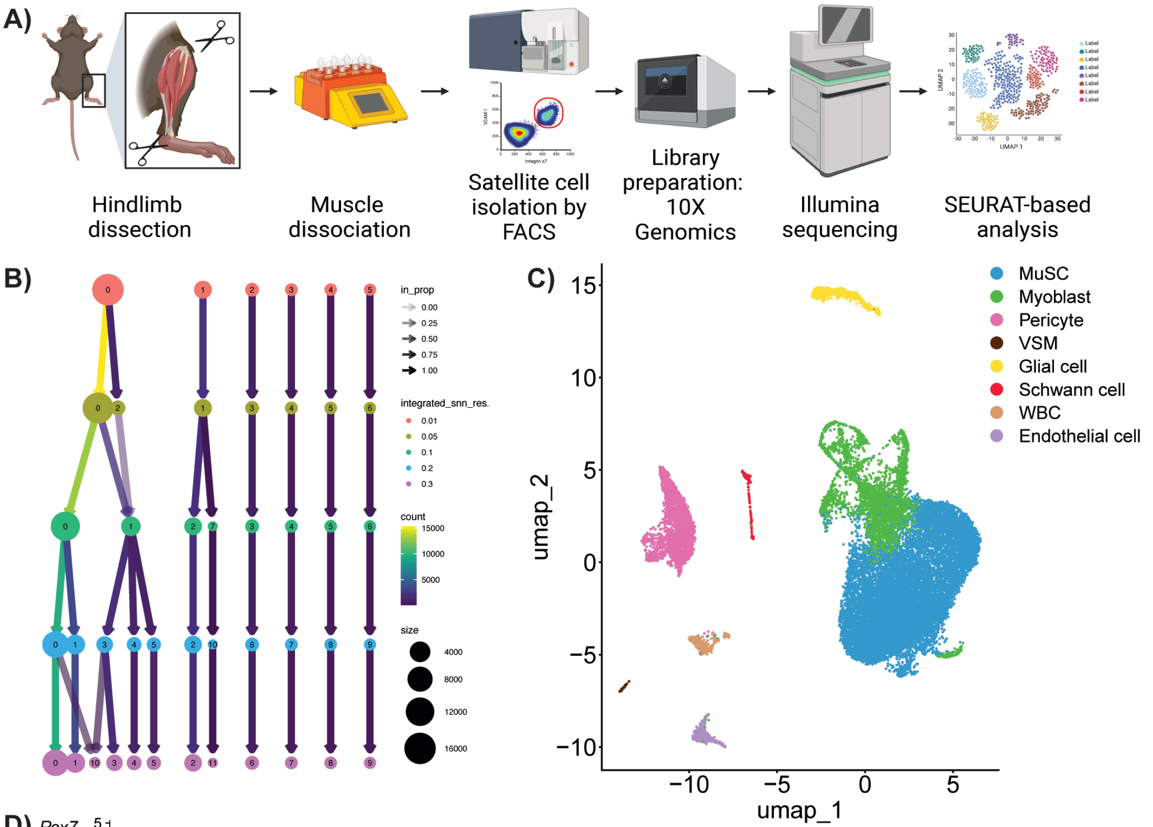


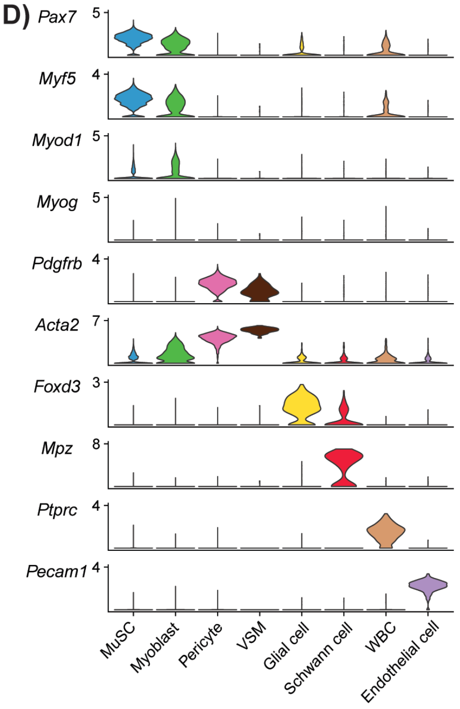


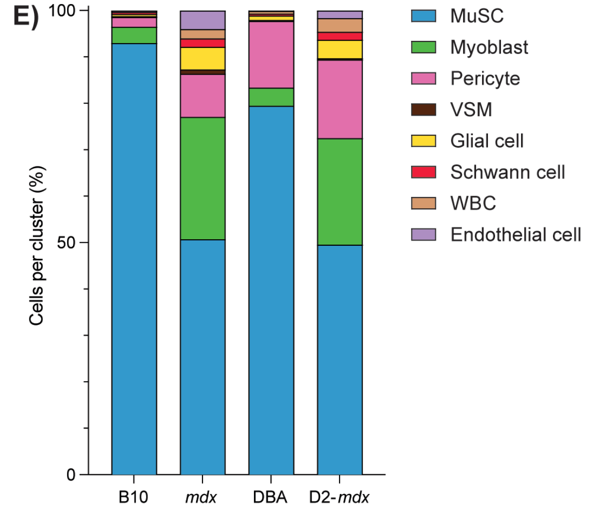


F)

| | MuSC | Myoblast | Pericyte | VSM | Glial cell | Schwann cell | WBC | Endothelial cell |
|---|---|---|---|---|---|---|---|---|
| **B10** | 92.97 | 3.50 | 2.13 | 0.15 | 0.49 | 0.43 | 0.04 | 0.28 |
| ***mdx*** | 50.72 | 26.39 | 9.28 | 0.92 | 4.88 | 1.82 | 2.02 | 3.98 |
| **DBA** | 79.46 | 3.93 | 14.36 | 0.21 | 0.94 | 0.40 | 0.61 | 0.09 |
| **D2-*mdx*** | 49.58 | 22.91 | 16.93 | 0.29 | 3.97 | 1.73 | 2.96 | 1.63 |

**Supplemental Figure 1. A)** Workflow schematic of scRNAseq, from hindlimb dissections to SEURAT-based analysis (created with BioRender.com). **B)** Clustree plot of all captured cells showing changes in clustering based on resolution 0.1. **C)** UMAP plot showing the grouping of all captured cells into eight clusters, of which the MuSCs (blue) and myoblast clusters (green) were used for further analysis. **D)** Violin plots of highly expressed markers used to define the identity of each cluster. **E)** Percentage of cells in each cluster, summarized in **F)**, showing an increase in the proportion of myoblasts and decrease in MuSCs in *mdx* and D2-*mdx* compared to their respective controls.

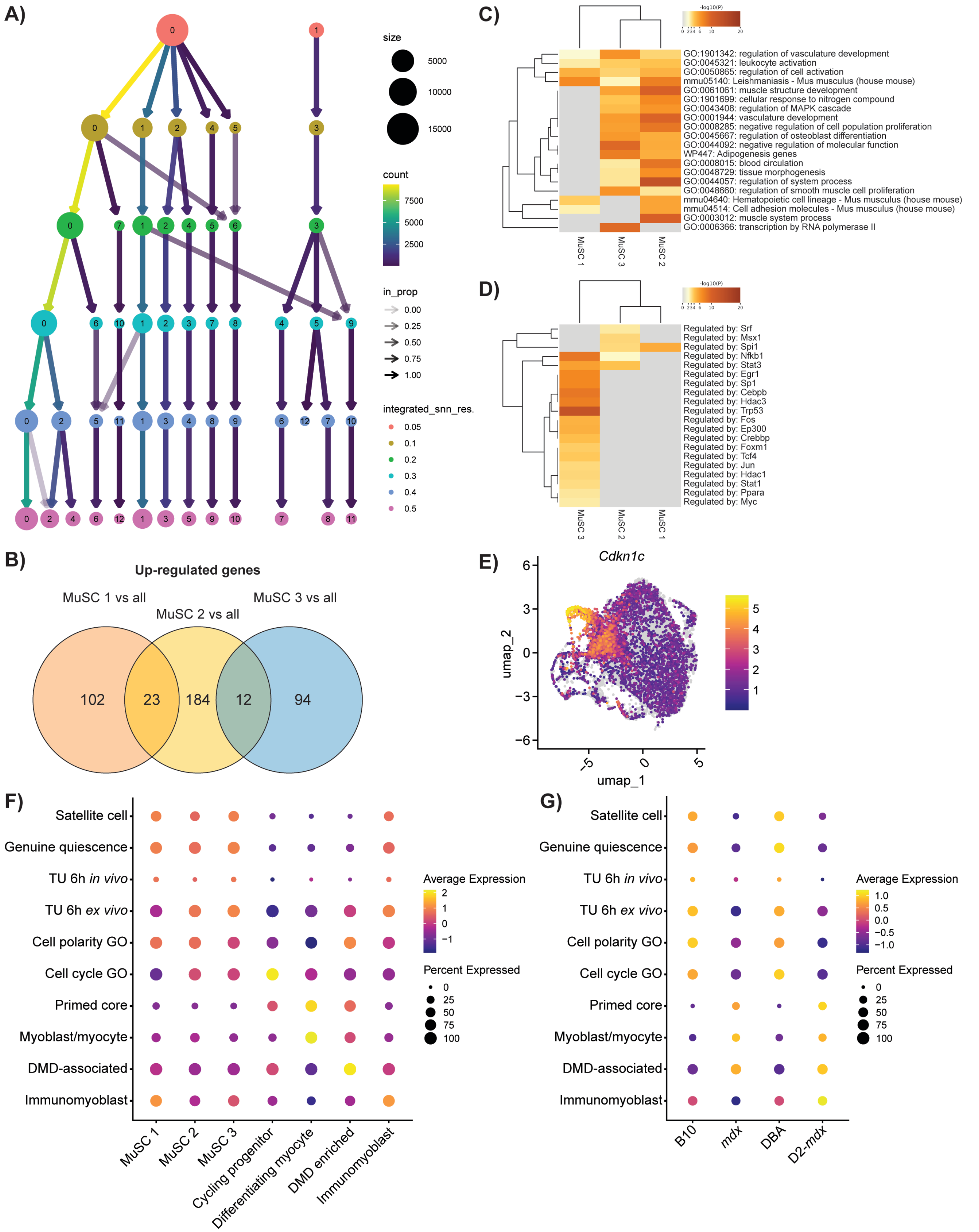


**Supplemental Figure 2. A)** Clustree plot of all myogenic cells showing changes in clustering based on resolution, with a resolution of 0.2 being used for further analyses. **B)** Venn diagram of upregulated genes (log2FC > 1 and p-value < 0.05) from muscle stem cell (MuSC) clusters 1, 2, and 3 versus all clusters showing overlap between MuSC 2 with MuSC 1 and MuSC 3 clusters, and no overlap between MuSC 1 and 3 clusters. **C)** Heatmap representing enriched terms, coloured by p-value, from genes upregulated in MuSC 1-3 clusters. **D)** Transcriptional regulatory relationships analysis (TRRUST) of genes upregulated in MuSC clusters 1-3 indicating enrichment of genes regulated by transcription factors associated with satellite cell activation in the MuSC 3 cluster. **E)** Feature plot of *Cdkn1c* showing increased expression in areas representing DMD enriched and differentiating myocyte clusters. **F)** Dot plot of expression of previously established scores by cluster type to validate identity, showing enrichment of the satellite cell, genuine quiescence, TU 6h *in vivo*, TU 6h *ex vivo*, and cell polarity GO scores in the MuSC 1-3 clusters and enrichment in primed core and myoblast/myocytes scores in cycling progenitor and differentiating myocyte clusters[1, 2, 3, 4, 5]. **G)** Dot plot of the previous scores displayed by strain, showing enrichment of satellite cell and genuine quiescence scores in healthy controls and enrichment in primed core, myoblast/myocyte, and DMD-associated scores in DMD models.

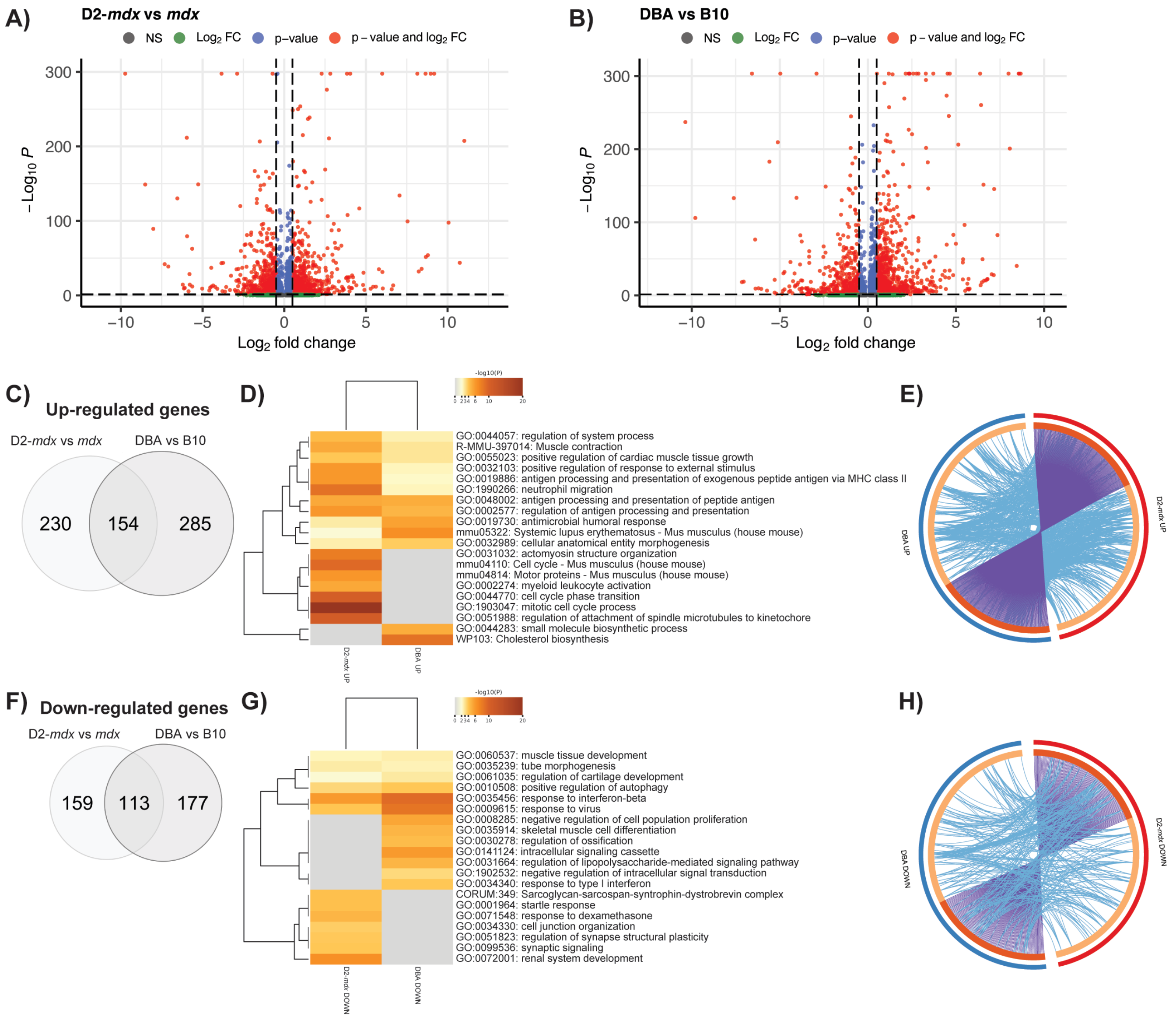


**Supplemental Figure 3.** Volcano plots of differentially expressed genes in **A)** D2-*mdx* versus *mdx* and **B)** DBA versus B10. **C, F)** Venn diagram of up-regulated (**C**) and down-regulated (**F**) genes from D2-*mdx* versus *mdx* (light grey) and DBA versus B10 (dark grey) and their overlap. **D, G)** Heatmaps of top 20 enriched terms, coloured by p-value, generated from these gene lists, showing minimal effect of strain background. **E, H)** Circos plots representing overlap between up-regulated (**E**) and down-regulated (**H**) gene lists. Outer circle represents the gene list for DBA (blue) and D2-*mdx* (red). Inner circle represents gene lists, where hits are arranged along the arc. Genes that hit multiple lists are colored in dark orange, and genes unique to a list are shown in light orange. Purple curves link identical genes between gene lists. Blue curves link genes that belong to the same enriched ontology term.

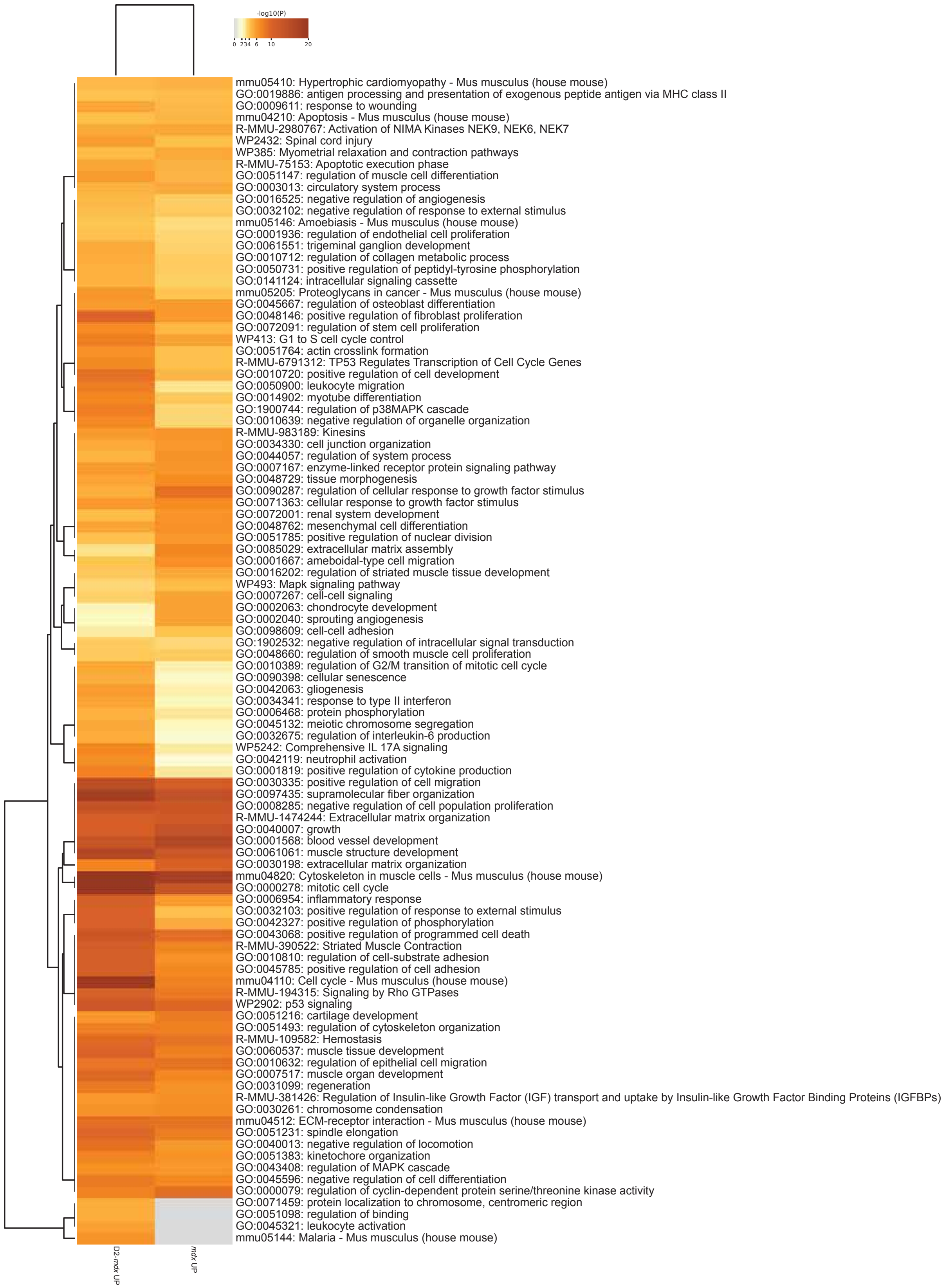


**Supplemental Figure 4.** Heatmap of top 100 enriched terms, coloured by p-value, generated from differentially expressed upregulated genes (log2FC > 1 and P < 0.05) in *mdx* vs B10 and D2-*mdx* vs DBA.

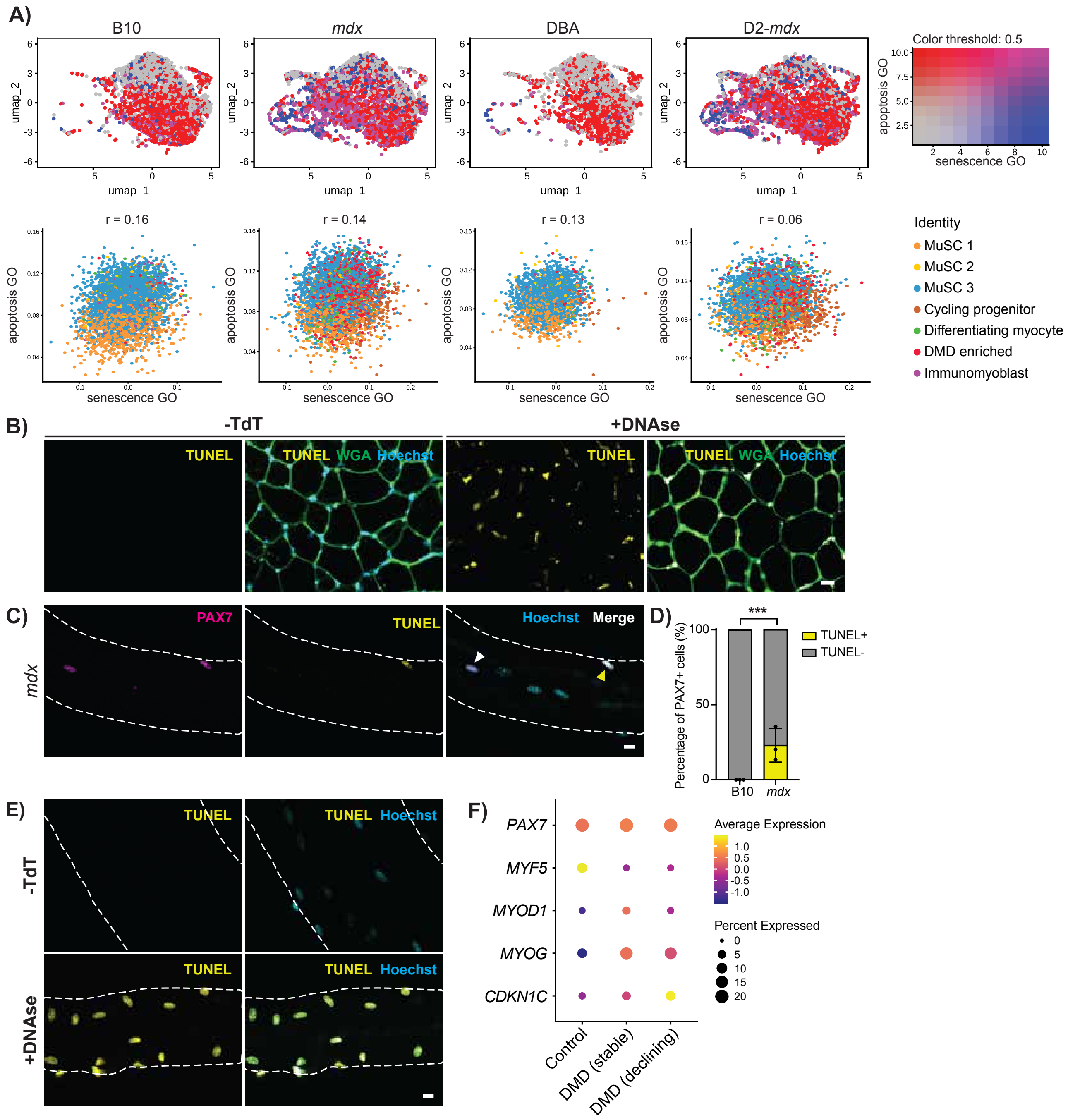


**Supplemental Figure 5. A)** Feature plots for each mouse strain overlapping the expression of the apoptosis (red) and senescence GO term (blue) scores, showing little overlap, and the scatterplots with the indicated Pearson correlation coefficients (r) for each respective strain, labelled by cell cluster identity, showing no correlation between the two scores. **B)** Controls for TUNEL staining on TA cross sections showing the negative control without terminal deoxynucleotide transferase (-TdT) and the positive control treated with DNAse. **C)** Representative IF labelling of an uninjured *mdx* EDL myofiber for PAX7 (magenta) and TUNEL (yellow) showing a nucleus positive for PAX7 only (white arrow) and a nucleus double positive for both PAX7 and TUNEL (yellow arrow). **D)** Quantification of the number of total PAX7+ nuclei that are positive for TUNEL, visualized as a percentage, showing their presence in *mdx* only (n = 3 biological replicates). **E)** Controls for TUNEL staining on *mdx* EDL myofibers showing the negative control without TdT and the positive control treated with DNAse. **F)** Dot plot of myogenic factors and *CDKN1C* expression in human satellite cells generated from a published single nuclei transcriptomic dataset from DMD patients and age-matched controls[6]. ***P < 0.001 (two-tailed unpaired t-test, n = 3 biological replicates per strain). Data are expressed as mean ± SD. Scalebars = 20 μm (**B**), 10 μm (**C**, **E**).

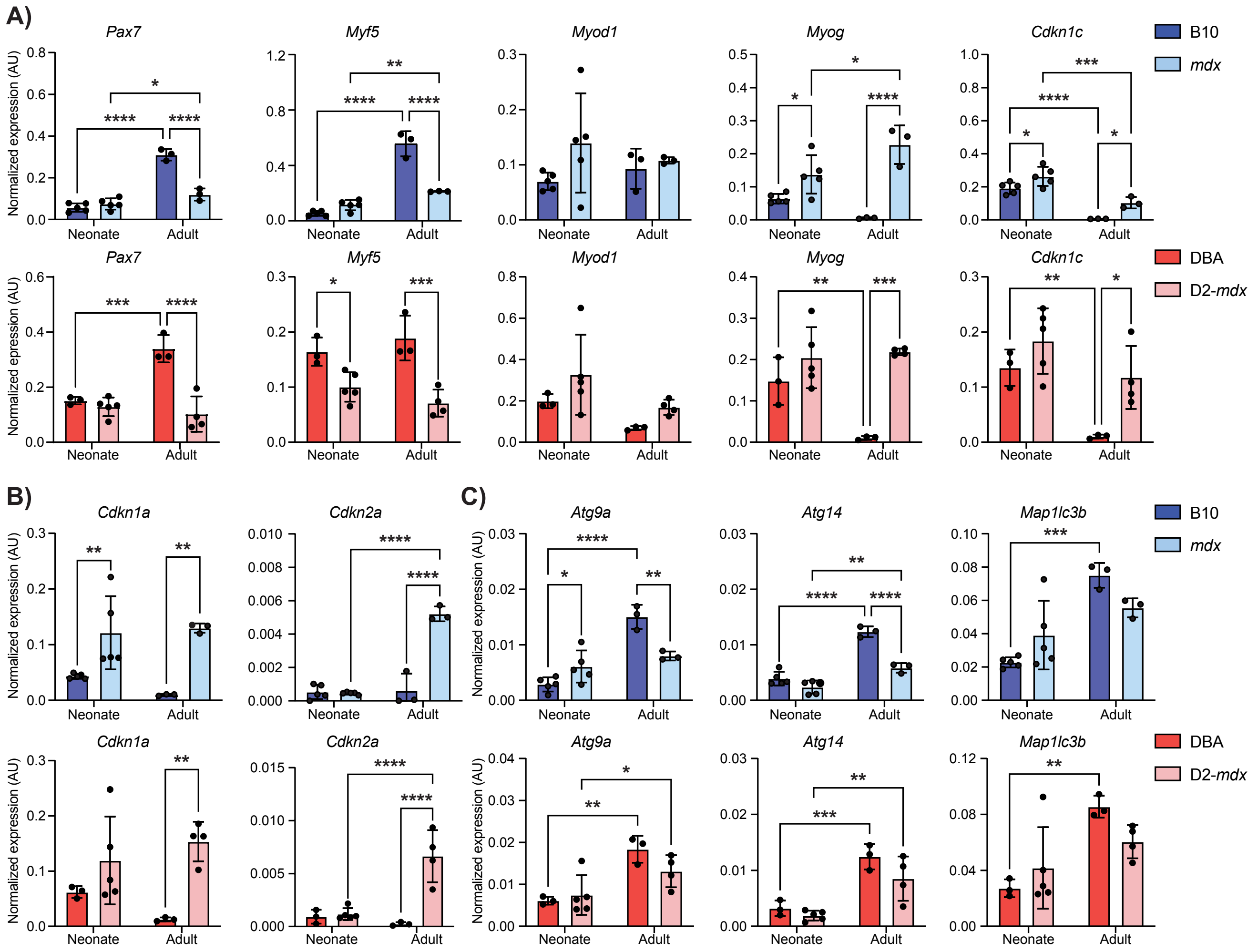


**Supplemental Figure 6.** Digital PCR quantification of **A)** myogenic regulatory factors, **B)** senescence-associated genes and **C)** autophagy-associated genes from satellite cells prospectively isolated from neonate (10-14 days-old) and adult (three months-old) B10, *mdx*, DBA and D2-*mdx* mice showing some early changes in DMD models as compared to wildtype controls. *P < 0.05, **P < 0.01, ***P < 0.001, ****P < 0.0001 (two-way analysis of variance). Data are expressed as mean ± SD.

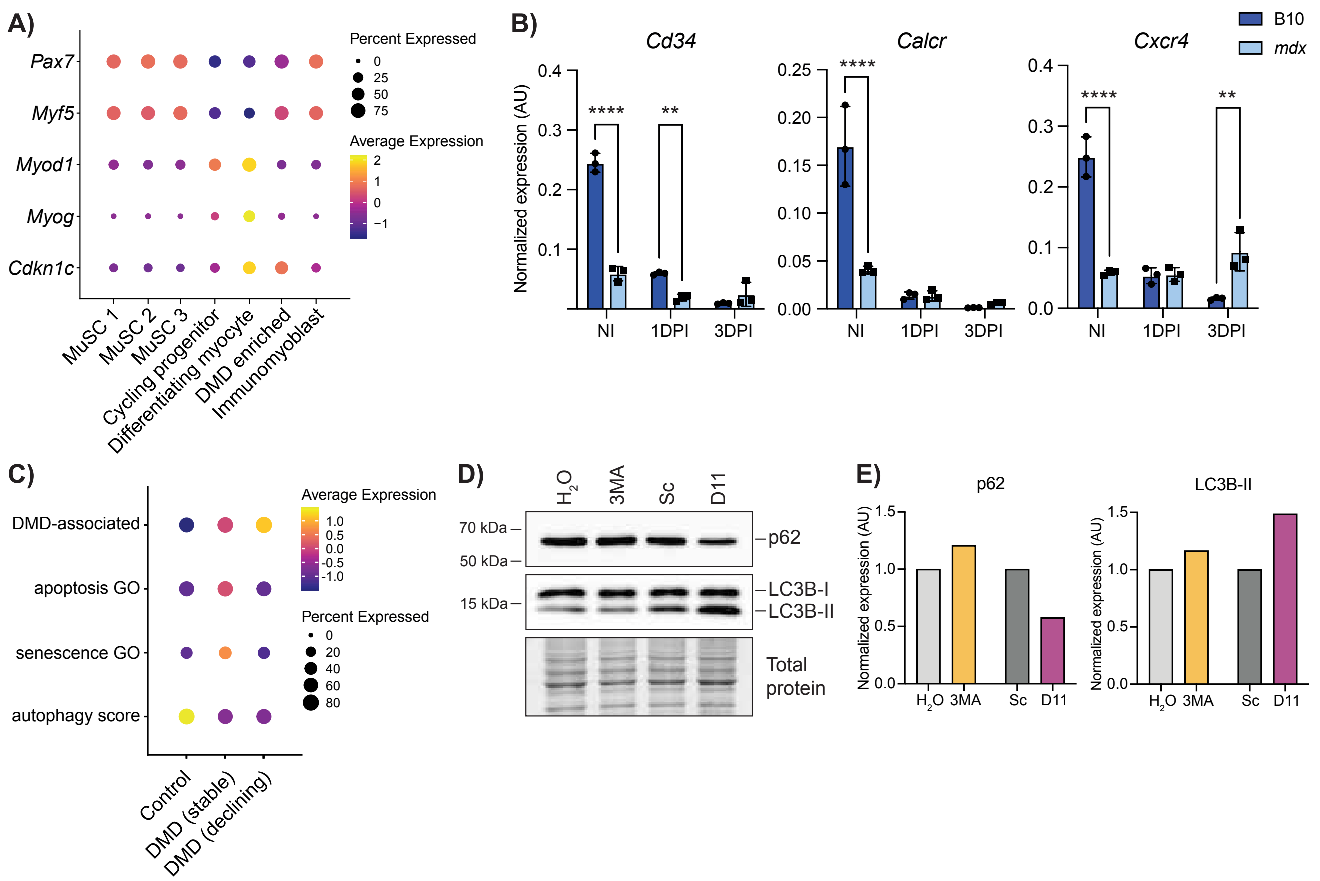


**Supplemental Figure 7. A)** Dot plot of myogenic factors and *Cdkn1c* expression in the scRNA clusters showing higher expression of satellite cell markers *Pax7* and lower expression of progenitor markers *Myod1* and *Myog* in the DMD enriched cluster compared to the cycling progenitor and differentiating myocyte clusters. **B)** Expression of quiescence-associated genes as measured by ddPCR in isolated satellite cells from NI, 1DPI and 3DPI muscle from B10 and *mdx* mice showing less expression in *mdx* at NI and 1DPI. **C)** Dot plot of expression DMD-associated, apoptosis GO, senescence GO, and autophagy scores in human satellite cells generated from a published single nuclei transcriptomic dataset from DMD patients and age-matched controls[6]. **D)** Western blot showing levels of p62 (top) and LC3B (middle) after 3MA and D11 treatments and their respective controls, as well as total protein levels (bottom). Expression of p62 is reduced, while LC3B-II is increased, following D11 treatment compared to Sc control. **E)** Quantification of p62 and LC3B-II expression levels, normalized to total protein, and presented as relative to respective controls. $^{**}P < 0.01$, $^{****}P < 0.0001$ (two-way analysis of variance). Data are expressed as mean ± SD.
